# Aberrant DNA repair is a vulnerability in histone H3.3-mutant brain tumors

**DOI:** 10.1101/2022.09.29.510093

**Authors:** Beatrice Rondinelli, Giulia Giacomini, Sandra Piquet, Odile Chevallier, Juliette Dabin, Siau-Kun Bai, Byungjin Kim, Robert Siddaway, Brian Raught, Etienne Coyaud, Chun-Min Shan, Robert J.D. Reid, Takenori Toda, Rodney Rothstein, Therese Wilhelm, Viviana Barra, Alexander Crane, Frank Dubois, Pratiti Bandopadhayay, Rameen Beroukhim, Valeria Naim, Songtao Jia, Cynthia Hawkins, Sophie E. Polo

## Abstract

Pediatric high-grade gliomas (pHGG) are devastating and incurable brain tumors with recurrent mutations in histone H3.3. These mutations promote oncogenesis by dysregulating gene expression through alterations of histone modifications. We identify aberrant DNA repair as an independent oncogenic mechanism, which fosters genome instability and tumor cell growth in H3.3 mutant pHGG, thus opening new therapeutic options. The two most frequent H3.3 mutations in pHGG, K27M and G34R, drive aberrant repair of replication-associated damage by non-homologous end joining (NHEJ). Aberrant NHEJ is mediated by the DNA repair enzyme Polynucleotide Kinase 3’-Phosphatase (PNKP), which shows increased association with mutant H3.3 at damaged replication forks. PNKP sustains the proliferation of cells bearing H3.3 mutations, thus conferring a molecular vulnerability, specific to mutant cells, with potential for therapeutic targeting.

## Introduction

Pediatric high-grade gliomas (pHGGs) are deadly brain tumors with less than ten percent survival within two years of diagnosis. Despite aggressive radio/chemotherapy regimens, pHGGs remain incurable and are the leading cause of cancer-related death in children (Pinto et al., 2021). The toxicity of existing therapies and the emergence of resistance hinder the efficacy of current therapeutic protocols and call for the development of alternative, targeted therapeutic strategies that exploit specific molecular features of cancer cells (Lin et al., 2022; Pinto et al., 2021). One such feature is exemplified by heterozygous, dominant point mutations in the H3F3A gene, which encodes the H3.3 histone variant. These recurrent mutations are active players in the oncogenic process in pHGG (Deshmukh et al., 2021), with three mutually exclusive mutations resulting in single amino-acid substitutions at Lysine 27 to Methionine (H3.3K27M), giving rise to midline tumors of the central nervous system, or at Glycine 34 to Arginine or Valine (H3.3G34R/V) in cerebrocortical tumors (Schwartzentruber et al., 2012; Sturm et al., 2012; Wu et al., 2012). These mutations perturb histone post-translational modifications (PTMs) in cis (Fang et al., 2018; Shi et al., 2018) or in trans (Fang et al., 2016; Lewis et al., 2013), thus interfering with genome-wide gene expression programs, stalling the differentiation of interneuron progenitor cells at different stages (Bjerke et al., 2013; Chen et al., 2020a; Filbin et al., 2018; Jessa et al., 2019) and fueling cell transformation (reviewed in (Deshmukh et al., 2021; Phillips et al., 2020; Sahu and Lu, 2022)).

Besides their effect on gene expression, pHGG H3.3 mutations also promote genome instability (Bočkaj et al., 2021; Fang et al., 2018; Haase et al., 2022; Pfister et al., 2014; Schwartzentruber et al., 2012; Sturm et al., 2012; Yadav et al., 2017), which is one of the major drivers of cellular transformation (Hanahan, 2022). In particular, H3.3 mutant pHGGs display increased copy number alterations and chromosomal rearrangements compared to wild-type (WT) H3.3 tumors (Bočkaj et al., 2021; Mackay et al., 2017; Schwartzentruber et al., 2012). Mechanistically, H3 G34 mutations in fission yeast and mammals and the non-pHGG H3.3 K36M mutation in human cells hijack the response to DNA damage (Fang et al., 2016, 2018; Lowe et al., 2021; Pfister et al., 2014; Yadav et al., 2017) by dominantly interfering with the DNA repair-promoting function of WT H3.3 (Juhász et al., 2018; Luijsterburg et al., 2016) or by downregulating DNA damage response genes (Haase et al., 2022). Moreover, K27M and G34 mutations alter H3.3 association with several DNA repair factors in human cells (Fang et al., 2018; Lim et al., 2017; Siddaway et al., 2022). These observations expand the role of mutated H3.3 proteins beyond their ability to dysregulate gene expression and pave the way for the identification of additional oncogenic functions. However, mechanistic understanding of the impact of pHGG H3.3 mutations on DNA repair and genome integrity in human cells is still lacking, and whether and how this function can be exploited for therapy remain underexplored.

## Results

### H3.3 mutants drive aberrant DNA repair in S phase

To study the impact of various H3.3 mutations on DNA repair in human cells in an isogenic context, we generated U2OS cell lines stably expressing SNAP-tagged WT or individual mutant H3.3 proteins (bearing K27M, G34R/V pHGG mutations, and G34W, K36M non-pHGG mutations) in an H3.3 WT background (Figure S1A). The engineered cell lines have comparable expression of the different H3.3-SNAP proteins (Figure S1B-C). They also recapitulate some of the histone PTM changes (H3K27me3 and H3K36me3) and the mutant H3.3 to WT H3 ratio that characterize H3.3 mutant pHGGs (Fang et al., 2016; Lewis et al., 2013) (Figure S1B-C). In this system, we analyzed the focal accumulation of DNA repair factors upon treatment with different genotoxic agents. Similar to the positive control H3.3K36M (Fang et al., 2016), upon replication stress, two pHGG mutants, H3.3K27M and G34R, showed impaired foci formation of the RAD51 recombinase (RAD51) (Figure 1A), and of Fanconi Anemia Complementation Group D2 (FANCD2)(Figure S2A), both of which are involved in pathways that preferentially repair replication-associated DNA damage (Scully et al., 2019). We evaluated a possible compensatory activation of the non-homologous end joining (NHEJ) repair pathway in the same cell lines and observed increased foci formation of TP53-binding protein 1 (53BP1) (Figure 1B), a positive regulator of NHEJ (Zhao et al., 2020). We noticed a similar increase of 53BP1 foci in cells expressing the non-pHGG G34W oncomutant (Figure 1B). The altered recruitment of repair factors was detectable upon interference with replication fork (RF) progression (Berti et al., 2020) by camptothecin (CPT), hydroxyurea (HU) or mitomycin C (MMC), but not upon treatment with the radiomimetic agent bleomycin, which triggers DNA damage throughout the cell cycle (Figure 1A-B, Figure S2A-C), indicating that H3.3K27M and G34R mutants skew the repair of RF-associated DNA lesions. Importantly, the observed defect is not due to a different cell cycle distribution of cells expressing mutant H3.3 (Figure S2D), to differential signaling of DNA damage, as shown by comparable levels of gH2AX (Figure S2E), nor to differential abundances of repair proteins (Figure S2A and S2F). To functionally analyze NHEJ activity, we exploited the random plasmid integration assay, which confirmed increased NHEJ activity in cells expressing the pHGG mutants H3.3 K27M and G34R, but not G34V (Figure 1C, Figure S2G). The G34R and G34V histone H3 mutants also play opposing roles in DNA damage repair in fission yeast (Lowe et al., 2021; Yadav et al., 2017). This prompted us to exploit this model system to assess the possible conservation of increased, aberrant NHEJ activity in cells expressing pHGG mutants. Fission yeast contains three identical histone H3 genes and we introduced the K27M, G34R and G34V mutations into one of the H3 genes, while leaving the other two intact. Proliferation analyses showed that deletion of the core NHEJ factor Xrc4 (*xrc4Δ*) sensitized H3 WT and G34V strains to CPT damage, consistent with a protective role of NHEJ in these strains, which was not observed in K27M and G34R strains (Figure 1D). *xrc4Δ* even rescued the CPT sensitivity of G34R yeast cells, as observed both in proliferation and serial dilution plating assays, indicating that aberrant NHEJ drives the sensitivity of this strain to CPT.

**Figure 1.**
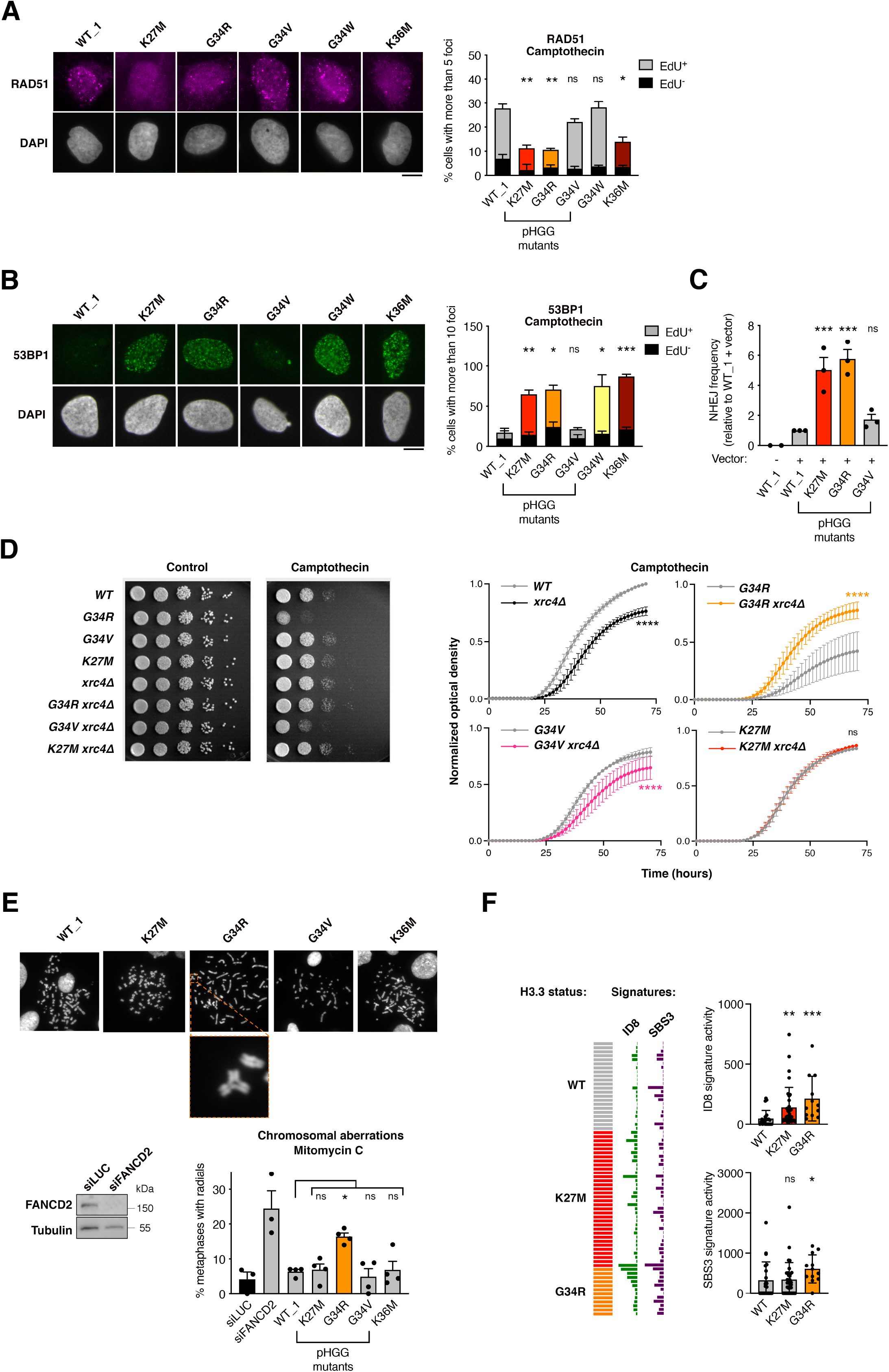
H3.3K27M and G34R pHGG mutations hijack DNA repair in S phase and harbor genomic instability features of aberrant NHEJ. (A-B) Analysis of RAD51 (A) and 53BP1 (B) repair foci by immunofluorescence in U2OS cells stably expressing wild-type H3.3 (WT_1) or the indicated mutants and treated with camptothecin (3 h, 0.1 μM). Representative images of repair foci in EdU^+^ cells are shown. Bar graphs depict the percentage of EdU^+^ and EdU^-^ cells harboring more than 5 (RAD51) or 10 (53BP1) foci. Mean ± SEM from two to three independent experiments, with n>117 per sample for each experiment. (C) Analysis of NHEJ activity by random plasmid integration assay in U2OS cells stably expressing wild-type (WT_1) or mutant H3.3. Vector –, negative untransfected control. (D) Serial dilution analyses and proliferation curves of *S. pombe* strains expressing wild-type or mutant H3, depleted for the core NHEJ factor Xrc4 (*xrc4*Δ) and grown in standard growth medium (control) or in the presence of camptothecin (10 μM). (E) Scoring of radials in metaphase spreads of U2OS cells stably expressing wildtype (WT_1) or mutant H3.3 and treated with Mitomycin C (24 h, 25 ng/mL). Cells transfected with siRNA against FANCD2 (siFANCD2) are used as positive control (siLUC, siLuciferase, control). A representative example of a radial chromosome is shown in the inset. The western blot shows siRNA efficiency (Tubulin, loading control). Mean ± SEM from three independent experiments, with n>49 per sample for each experiment. (F) Oncoprint representation of DNA repair-driven mutational signatures (ID8, indels 8; SBS3, single-base substitutions 3) in whole-genome sequences of pre-treatment, TP53-mutated, primary pHGG samples harboring wild-type H3.3 (WT), H3.3K27M or G34R. Statistical significance is calculated by two-way ANOVA (A, B), one-way Anova (C, E), non-linear regression analysis with a polynomial quadratic model (D) or the non-parametric Kruskal-Wallis test (F). *: p< 0.05; **: p< 0.01; ***: p< 0.001; ns: p> 0.05. Scale bars, 10 μm. See also Figures S1 and S2.

To test whether aberrant DNA repair in H3.3K27M and G34R cells is associated with genome instability in human cells, we examined the occurrence of radials, i.e. chromosomal aberrations that derive from mis-joining of broken chromatids through aberrant NHEJ (Zhao et al., 2020). We observed a marked accumulation of radials in H3.3G34R U2OS cells upon MMC treatment (Figure 1E). To conclusively link the aberrant DNA repair in H3.3 K27M and G34R cells to genome instability onset in a glioma context, we analyzed whole genome-sequencing data from a panel of pre-treatment, TP53-mutant primary pHGGs for the presence of mutational signatures generated by defects in DNA double-strand break repair pathways (Alexandrov et al., 2020). Both H3.3 K27M and G34R pHGGs presented higher levels of mutational signatures deriving from aberrant NHEJ (ID8) compared to WT H3.3 pHGGs, and H3.3 G34R also displayed a stronger signature activity of defective homologous recombination (SBS3) (Figure 1F). Collectively, our data support a model where the pHGG mutations H3.3K27M and G34R skew the repair of S phase DNA damage towards aberrant NHEJ, thus sustaining a specific pattern of genome instability.

### Aberrant NHEJ is independent of H3K27/K36me3 loss

To test whether pHGG H3.3 mutants skew the repair of S phase damage through gain- or loss-of-function mechanisms, we first evaluated the impact of siRNA-mediated depletion of H3.3 on RAD51 and 53BP1 focus formation in CPT-damaged U2OS cells. Depletion of H3.3 did not affect the proportion of cells in S phase (Figure S3A) nor gH2A.X induction in response to CPT (Figure S3B). Contrary to H3.3 mutations, H3.3 loss did not alter RAD51 and 53BP1 focus formation in response to CPT (Figure 2A), despite an increase in 53BP1 nuclear levels (Figure S3C). Thus, H3.3 K27M and G34R mutations do not phenocopy H3.3 loss but rather confer a new function to histone H3.3 upon CPT-induced damage, corroborating the gain-of-function hypothesis.

**Figure 2.**
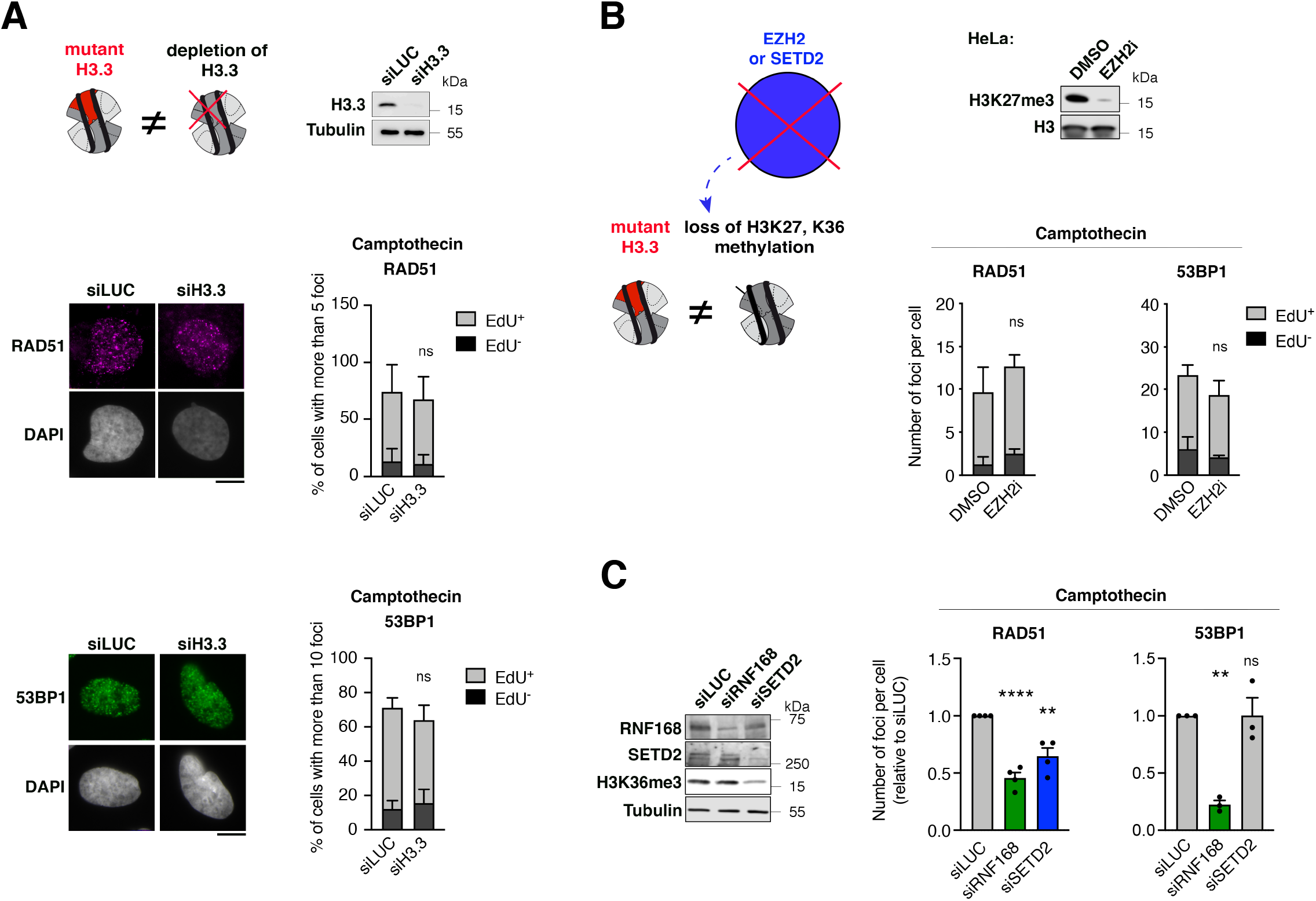
pHGG H3.3 mutants hinder DNA repair through a gain-of-function mechanism independently of hypomethylation at Lysines 27 and 36 of histone H3. (A) Analysis of RAD51 and 53BP1 foci by immunofluorescence in U2OS cells transfected with siRNAs against Luciferase (siLUC, control) or H3.3 (siH3.3) and treated with camptothecin (3 h, 0.1 μM). The western blot shows siRNA efficiency (Tubulin, loading control). Representative images of RAD51 and 53BP1 foci in EdU^+^ cells are shown. Bar graphs depict the percentage of EdU^+^ and EdU^-^ cells harboring more than 5 (RAD51) or 10 (53BP1) foci. Mean ± SEM from two independent experiments, with n>113 per sample for each experiment. (B) Analysis of RAD51 and 53BP1 foci by immunofluorescence in HeLa cells treated with DMSO or the EZH2 inhibitor GSK126 (EZH2i, 72 h, 1 μM) and damaged with camptothecin (3 h, 0.1 μM). The western blot shows the efficiency of EZH2 inhibition by analyzing H3K27me3 levels. Bar graphs depict the number of RAD51 or 53BP1 foci per cell in EdU^+^ and EdU^-^ cell populations. Mean ± SEM from three independent experiments, with n>125 per sample for each experiment. (C) Analysis of RAD51 and 53BP1 foci by immunofluorescence in U2OS cells transfected with the indicated siRNAs (siLUC, negative control; siRNF168, positive control known to inhibit both RAD51 and 53BP1 foci formation) and treated with camptothecin (3 h, 0.1 μM). siRNA efficiencies and H3K36me3 levels are analyzed by western blot (Tubulin, loading control). Bar graphs depict the number of RAD51 or 53BP1 foci per cell. Mean ±SEM from three independent experiments, with n>131 for each experiment. Statistical significance is calculated by two-way ANOVA (A, B) or one-way ANOVA (C). *: p< 0.05; **: p< 0.01; ***: p< 0.001; ns: p> 0.05. Scale bars, 10 μm. See also Figure S3.

Dysregulation of gene expression programs by H3.3 K27M and G34R mutations is mediated by reduced trimethylation at Lysines 27 (Lewis et al., 2013; Venneti et al., 2013) and 36 (Bjerke et al., 2013; Fang et al., 2018) of histone H3 (H3K27me3 and H3K36me3), respectively. To study whether the observed DNA repair defect is mediated by analogous perturbations of histone PTMs, we reduced H3K27me3 and H3K36me3 by inhibiting or depleting the corresponding lysine methyltransferases. The H3K27 methyltransferase Enhancer of Zeste 2 (EZH2) is endogenously inhibited in U2OS cells (Jain et al., 2019; Ragazzini et al., 2019), thus preventing further reduction of H3K27me3 upon expression of H3.3 K27M (Figure S1B and S3D). Yet, aberrant DNA repair is still observed in H3.3 K27M U2OS cells (Figure 1A-B), arguing against a contribution of H3K27me3 reduction to this repair defect. We confirmed this hypothesis in HeLa cells by chemical inhibition of EZH2 (EZH2i) (Figure 2B), which did not recapitulate the DNA repair defect observed in H3.3 K27M-expressing U2OS cells (Figure 1A-B). Similarly, reducing H3K36me3 by depleting SET Domain Containing 2 (SETD2) did not result in increased 53BP1 foci formation but solely in the expected reduction of RAD51 foci formation (Aymard et al., 2014; Carvalho et al., 2014; Pfister et al., 2014) in CPT-treated U2OS cells (Figure 2C). Together, these experiments demonstrate that H3.3 K27M and G34R mutants skew DNA repair towards NHEJ in S phase by conferring a gain-of-function to histone H3.3, independently of hypomethylation of H3K27 and K36.

### Mutant H3.3 histones are deposited at damaged forks

The impact of H3.3 mutations on the repair of S phase DNA damage and the reported *de novo* deposition of H3.3 at sites of DNA damage (Adam et al., 2013; Juhász et al., 2018; Luijsterburg et al., 2016) prompted us to investigate whether WT and pHGG H3.3 mutant proteins were *de novo* deposited at damaged RFs. First, we exploited fluorescent labeling of SNAP-tagged, newly synthesized histone H3.3 in a model of RF blockage, where stably integrated Lac operon (LacO) arrays generate an obstacle to DNA polymerase progression when bound by the Lac repressor (LacR) (Figure 3A) (Beuzer et al., 2014). Upon RF blockage, monitored by gH2AX accumulation (Figure S3E), we found a local enrichment of newly synthesized H3.3 on the LacO array specifically in S phase cells (Figure 3A), revealing a previously uncharacterized *de novo* deposition of H3.3 at sites of a replication block. To further study the deposition of newly synthesized H3.3 proteins in both WT and mutant cells, we setup a novel imaging-based method, SNAP-PLA (Proximity Ligation Assay), that measures by PLA (Söderberg et al., 2006) the colocalization between biotin-labeled, newly synthesized SNAP-tagged histones and gH2AX at CPT-damaged RFs (Figure 3B). Thus, we could detect *de novo* deposition of WT H3.3 specifically in CPT-damaged, S phase cells (Figure 3B and Figure S3F), recapitulating data obtained in the LacO system (Figure 3A) and validating the SNAP-PLA approach. Moreover, we detected *de novo* deposition of H3.3 K27M and G34R mutant proteins at damaged RFs, which was comparable to that of WT H3.3, while the *de novo* deposition H3.3 G34V was significantly reduced (Figure 3B). We validated these findings through the proteomic-based isolation of proteins on nascent DNA (iPOND) upon RF damage with CPT. The enrichment of SNAP-tagged H3.3 at CPT-damaged RFs was enhanced in S phase synchronized cells, supporting an S phase-specific deposition of H3.3 at damaged RFs (Figure S3G). Using this method, we confirmed the deposition of H3.3 K27M and G34R mutant proteins at CPT-damaged RFs (Figure 3C). The deposition of H3.3 G34V was detectable in this assay but with somehow reduced levels compared to WT H3.3 (Figure 3C). These findings unravel a local function of H3.3 in RF protection and repair and suggest that the deposition of pHGG H3.3 mutant proteins may locally affect the chromatin landscape and/or the recruitment of repair factors at damaged RFs, ultimately skewing fork repair.

**Figure 3.**
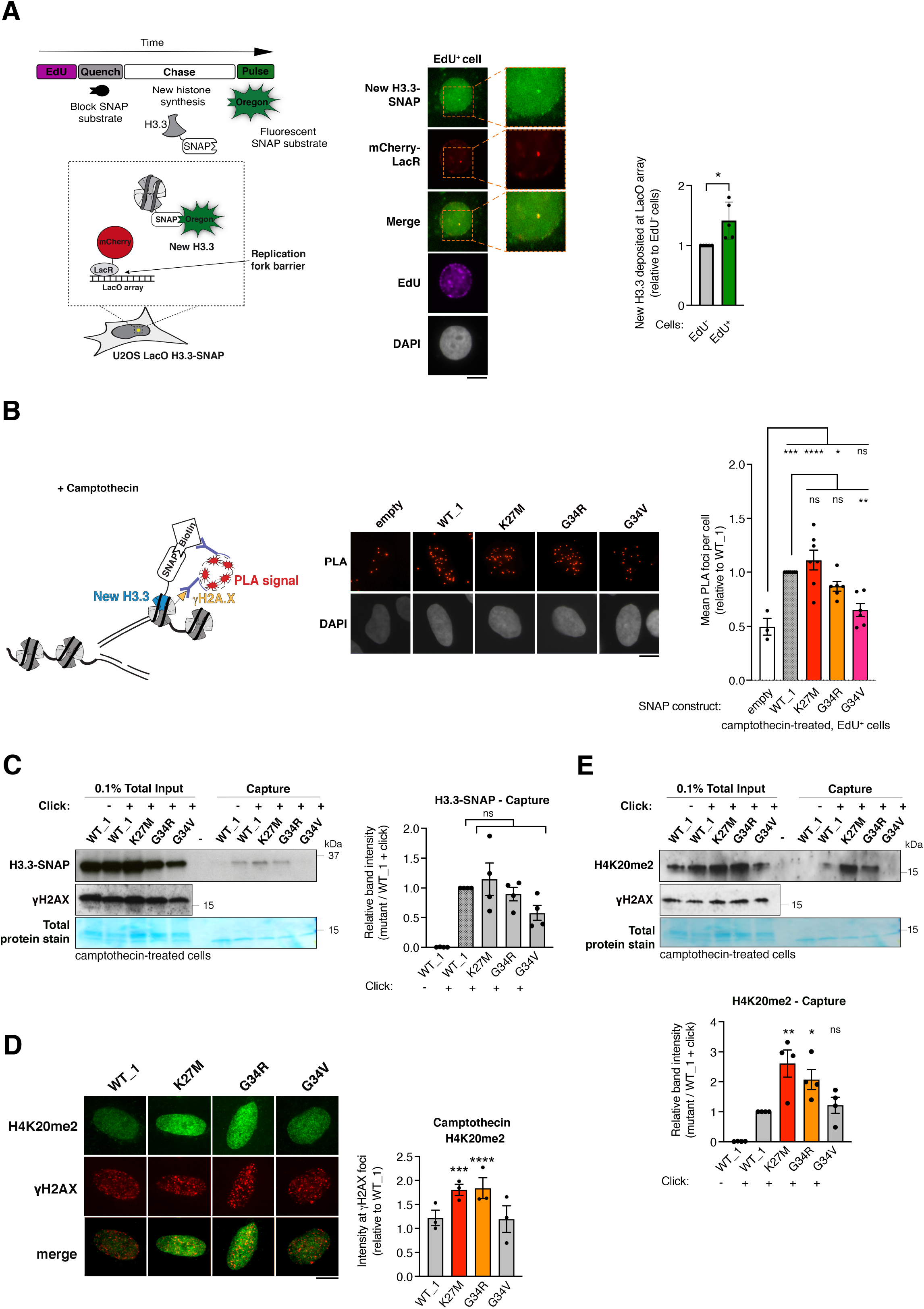
H3.3K27M and G34R pHGG mutants are *de novo* deposited and associated with increased H4K20me2 at damaged replication forks. (A) Scheme of the assay to monitor *de novo* deposition of wild-type H3.3-SNAP at the LacR-occupied LacO array fork barrier in U2OS LacO cells stably expressing SNAP-tagged H3.3 and transfected with mCherry-LacR. Images of a representative cell and 2.5x zoom on the LacO array. Quantification of new H3.3-SNAP accumulation (cells presenting an enrichment) at LacR-occupied LacO array in EdU^+^ and EdU^-^ cells. Mean ± SEM from five independent experiments, with n>20 per sample for each experiment. (B) Schematic representation of the SNAP-PLA assay to visualize the colocalization of gH2A.X with newly synthesized SNAP-tagged H3.3 (labeled with biotin) at RFs damaged with camptothecin (3 h, 0.1 μM). Representative images and quantification of SNAP-PLA colocalization foci between new H3.3 and gH2A.X in EdU^+^ U2OS cells stably expressing wild-type (WT_1) or mutant H3.3-SNAP, or SNAP tag only as a control (empty). Mean ± SEM from up to seven independent experiments, with n>130 per sample for each experiment. (C) Western blot analysis of input and capture samples from iPOND experiments performed in U2OS cells expressing wild-type (WT_1) or mutant H3.3-SNAP, synchronized in S phase and damaged with camptothecin (1 h, 1 μM). Click -, negative control (no biotin). Total protein stain shows the position of the streptavidin monomer, detectable at similar levels in all capture samples. Bar graphs depict H3.3-SNAP band intensity in capture samples relative to WT_1. Mean ± SEM from four independent experiments. (D) Immunofluorescence analysis of H4K20me2 levels at gH2A.X foci in U2OS cells stably expressing wild-type (WT_1) or mutant H3.3 and treated with camptothecin (3 h, 0.1 μM). Quantification of H4K20me2 intensity relative to WT_1. Mean ± SEM from three independent experiments, with n>17 per sample for each experiment. (E) Western blot analysis of input and capture samples from iPOND experiments performed in U2OS cells expressing SNAP-tagged wild-type (WT_1) or mutant H3.3, synchronized in S phase and damaged with camptothecin (1 h, 1 μM). Click -, negative control (no biotin). Bar graphs depict H4K20me2 band intensity in capture samples relative to WT_1. Mean ± SEM from three independent experiments. Statistical significance is calculated by unpaired t-test (A), one-way ANOVA (B-E). *: p< 0.05; **: p< 0.01; ***: p< 0.001; ns: p> 0.05. Scale bars, 10 μm. See also Figure S3.

To evaluate the impact of pHGG H3.3 mutant proteins on the chromatin landscape at damaged RFs, we analyzed by immunofluorescence the levels of H4K20me2, a histone mark known to drive 53BP1 foci formation on damaged chromatin (Botuyan et al., 2006). H4K20me2 nuclear levels were consistently higher in H3.3 K27M and G34R mutant cells (Figure S3H) including at CPT-damaged RFs marked by gH2A.X (Figure 3D). These findings are in line with the increased capacity of those cells to form 53BP1 foci (Figure 1B). Importantly, we also detected increased H4K20me2 at CPT-damaged RFs by iPOND in cells expressing H3.3 K27M and G34R compared to cells expressing WT H3.3 or the G34V mutant (Figure 3E). Together, these data establish that the deposition of H3.3 K27M and G34R mutant proteins at damaged RFs is associated with dysregulated histone PTMs that may, in turn, affect fork repair by impacting the recruitment of repair factors.

### PNKP associates with mutant H3.3

To identify DNA repair factors that preferentially associate with the H3.3K27M and G34R mutants, we employed proximity-dependent biotinylation (BioID) (Roux et al., 2012; Scott and Campos, 2020) in HEK293 human cells ectopically expressing WT, K27M or G34R H3.3 fused to the mutant BirA* biotin ligase, followed by mass spectrometry analysis (Figure 4A, Table S1 and (Siddaway et al., 2022)). Validating this approach, we detected the expected preferential association of EZH2 with the H3.3 K27M mutant (Lewis et al., 2013), while Nuclear Receptor Binding SET Domain Protein 1 (NSD1), responsible for H3K36 mono and dimethylation, showed reduced association to H3.3 G34R, in accordance with the reduced methylation of H3K36 by NSD1 in the presence of G34R (Fang et al., 2018) (Figure 4A). Among the DNA repair enzymes that preferentially associated with both K27M and G34R compared to WT H3.3, we focused our attention on the DNA end processing enzyme Polynucleotide Kinase 3’-Phosphatase (PNKP, Figure 4A), which contributes to NHEJ by transferring a phosphate group between broken DNA ends before ligation (Dumitrache and McKinnon, 2017). In addition, PNKP was identified as an H3.3G34R interactor in a previous study (Lim et al., 2017) and plays a central role in neurodevelopment (Shen et al., 2010). PNKP total levels were not increased in cells expressing H3.3 K27M or G34R (Figure S4A). However, by iPOND, we observed increased binding of PNKP to CPT-damaged RFs in H3.3 K27M and G34R cells (Figure 4B), further substantiating the preferential association of this DNA repair enzyme with both H3.3 mutant proteins.

**Figure 4.**
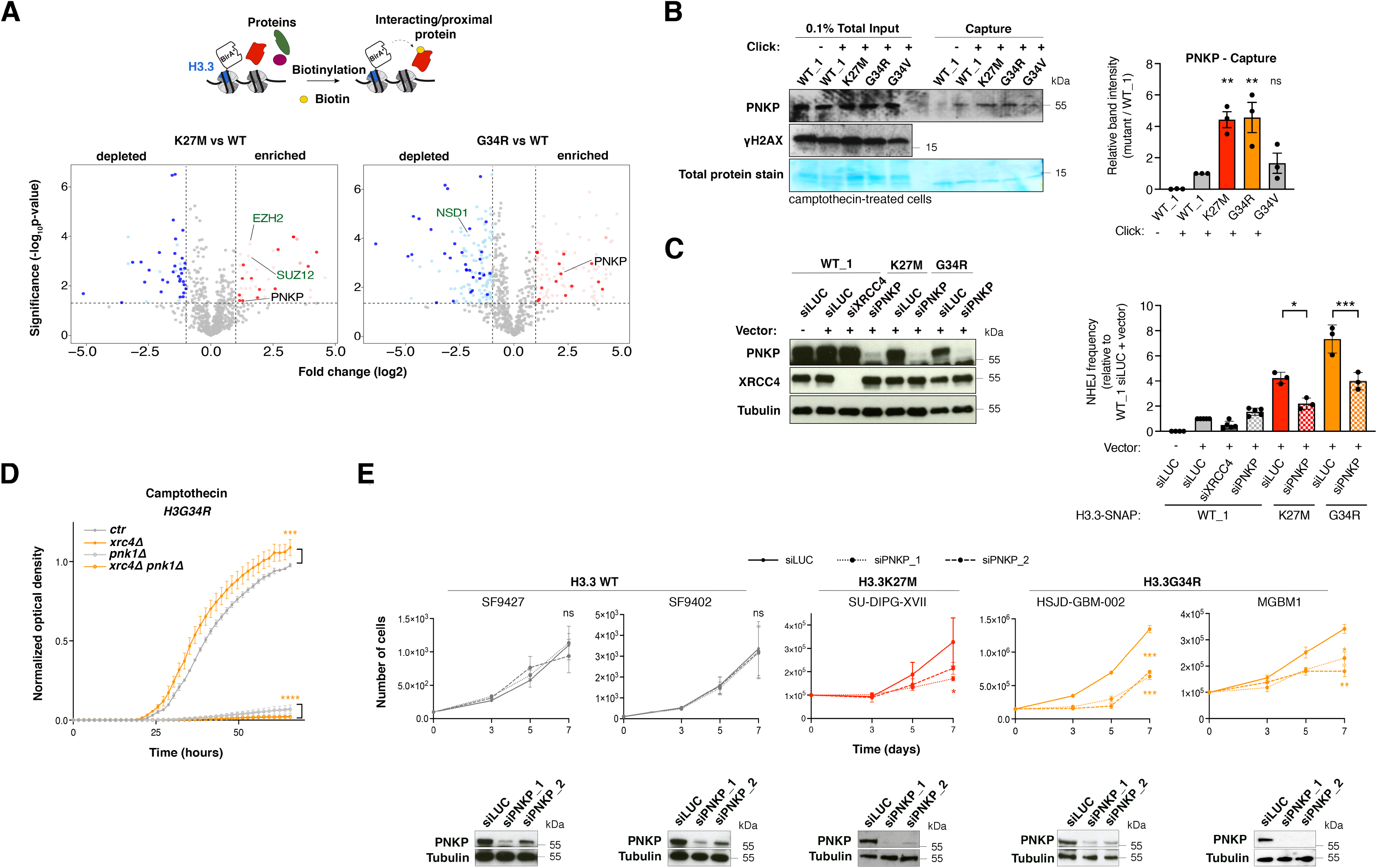
Abnormal PNKP function in H3.3 K27M and G34R mutant cells mediates aberrant NHEJ and represents a therapeutic target in pediatric pHGG. (A) Identification of proteins associated with wild-type (WT) and mutant H3.3 (K27M, G34R) by proximitydependent biotinylation (BioID) in HEK293 cells expressing BirA*-tagged H3.3 proteins. Volcano plots show interactors enriched (red) or depleted (blue) in H3.3K27M (left) or G34R (right) samples compared to WT H3.3 sample, with each dot representing an interactor. Significant interactors whose log2 fold change is > 1 and whose p-value is < 0.05 (−log_10_ p-value > 1.30) are highlighted in colors and common interactors between H3.3K27M and G34R are shown in dark colors. Positive controls are depicted in green. (B) Western blot analysis of input and capture samples from iPOND experiments performed in U2OS cells expressing wild-type (WT_1) or mutant H3.3-SNAP, synchronized in S phase and damaged with camptothecin (1 h, 1 μM). Click -, negative control (no biotin). Total protein stain shows the position of the streptavidin monomer, detectable at similar levels in all capture samples. Bar graphs depict PNKP band intensity in capture samples relative to WT_1. Mean ± SEM from three independent experiments. The representative experiment shown is the same as in Figure 3C. (C) Analysis of NHEJ activity by random plasmid integration assay in U2OS cells stably expressing wild-type (WT_1) or mutant H3.3 and transfected with siRNAs against Luciferase (siLUC, control) or PNKP (siPNKP). Samples siLUC -, siLUC + and siXRCC4 + are the same shown in Figure S2G graph. Western blot analysis shows siRNA efficiency (Tubulin, loading control). Vector -, negative untransfected control. (D) Proliferation curves of *S. pombe* strains expressing H3G34R, depleted for the core NHEJ factor Xrc4 (*xrc4*Δ) and for Pnk1 (*pnk1*Δ) and grown in the presence of camptothecin (5 μM). (E) Proliferation assays in patient-derived pHGG cell lines harboring wild-type or mutant H3.3 and transfected with siRNAs against Luciferase (siLUC, control) or PNKP (siPNKP_1 and siPNKP_2). The western blots show siRNA efficiencies (Tubulin, loading control). Statistical significance is calculated by one-way ANOVA (B, C), by non-linear regression analysis with a polynomial quadratic model (D, E). *: p< 0.05; **: p< 0.01; ***: p< 0.001; ns: p> 0.05. See also Figure S4 and Table S1.

### PNKP promotes aberrant repair in mutant cells

We hypothesized that the preferential association of PNKP with H3.3 K27M and G34R may drive the aberrant NHEJ observed in cells expressing these mutants. Thus, we measured NHEJ activity by random plasmid integration assay upon knockdown of PNKP in U2OS cells expressing WT or mutant H3.3. In contrast to the depletion of the core NHEJ factor XRCC4, PNKP knockdown did not affect NHEJ activity in WT H3.3 cells. However, knock-down of PNKP significantly reduced NHEJ in H3.3 K27M and G34R cells, showing that PNKP promotes aberrant NHEJ repair in these cells (Figure 4C). Similarly, in fission yeast, the rescue of CPT-sensitivity in the G34R strain by xrc4 deletion was not observed upon co-deletion of the PNKP ortholog pnk1, arguing that the aberrant xrc4-mediated NHEJ in a G34R background is dependent on pnk1 (Figure 4D). Together, these data demonstrate that PNKP drives aberrant NHEJ in H3.3 mutant cells and that this process is evolutionarily conserved.

### PNKP as a therapeutic target in pHGG

We next assessed the importance of aberrant PNKP function in H3.3 mutant cells. PNKP knockdown specifically impaired the growth of H3.3 K27M and H3.3G34R U2OS cells, but not of WT H3.3 or H3.3 G34V mutant cells (Figure S4B). By exploiting a panel of patient-derived glioma cells lines (Figure S4C) and two different siRNAs against PNKP, we corroborated the specific effect of PNKP knockdown on the proliferation of glioma cells harboring endogenous H3.3 K27M or G34R mutations (Figure 4E). PNKP knockdown significantly reduced the growth of all H3.3 mutant cell lines examined in contrast to H3.3 WT glioma cell lines, which were unaffected (Figure 4E). These data expand our findings to state-of-the-art pHGG cellular models and put forward PNKP as a potential therapeutic target in pHGG cells expressing specific H3.3 mutations.

## Discussion

By dissecting how H3.3 mutants skew DNA repair, we show that mutations in core chromatin components induce genome instability by dysregulating the recruitment of repair factors at damaged RF (Figure S4D). Aberrant repair of damaged RF leads to mitotic aberrations (Berti et al., 2020), which were recently reported in cells expressing the H3.3K27M mutant (Bočkaj et al., 2021) and is the likely cause of genome instability. The influence of co-occurring genetic alterations (Mackay et al., 2017) on this phenotype should be evaluated in future studies.

By identifying a new, local function of H3.3 in the repair of damaged RFs, the present study expands our knowledge about histone variant deposition at these sites (Kim et al., 2018; Xu et al., 2021). *De novo* H3.3 deposition may operate through a DNA synthesis-independent mechanism on the regressed arms of damaged RFs, which have been shown to be chromatinized (Schmid et al., 2018).

H3.3 K27M and G34R affect RF repair through a mechanism that is distinct from their interference with gene expression programs (Deshmukh et al., 2021; Haase et al., 2022; Phillips et al., 2020). The aberrant use of NHEJ in S phase indeed does not rely on H3K27me3 and H3K36me3 alterations, but may involve other PTM changes in mutant nucleosomes, possibly through the activity of histone modifying enzymes, which may in turn affect the binding of repair factors. For instance, the observed increase in H4K20me2 at RFs may explain enhanced 53BP1 foci formation and channeling of repair towards NHEJ.

The similarity of DNA repair phenotypes between H3.3 K27M and G34R mutant cells may indicate a common gain or loss of histone modifying activity on nucleosomes. The consistently stronger phenotype observed with the G34R mutant reflects the more efficient response to PNKP loss-of-function in H3.3 G34R mutant cells. In contrast, H3.3 G34R and G34V mutants display strikingly opposite DNA repair phenotypes, conserved from yeast to human, the molecular basis of which is still elusive. Perhaps the bulkier and positively charged arginine side chain of the G34R mutant protein causes a more drastic disruption of the H3.3 interactome. In addition, the G34V mutant is much rarer than G34R so maybe less oncogenic.

The K27M mutation is also found in the H3.1 histone variant (H3.1 K27M) in some pHGG and inhibits NHEJ in human fibroblasts (Zhang et al., 2018), which is opposite to the phenotype of H3.3 K27M in U2OS cells. Although a true comparison requires that the H3.1 K27M and H3.3 K27M phenotypes be studied in the same cellular background, differences in their DNA repair function can be anticipated since they show distinct distribution patterns in chromatin (Sarthy et al., 2020), present different co-occurring mutations (Hauser, 2021; Mackay et al., 2017) and clinical features in pHGG (Castel et al., 2015; Werbrouck et al., 2019).

Targeting DNA repair defects is the most developed approach so far to induce tumor cell killing through synthetic lethality, even if weakly exploited beyond BRCA-mutated tumors (Setton et al., 2021). By providing molecular understanding of the aberrant DNA repair capacities of H3.3 mutant pHGGs, we propose here that a synthetic interaction approach for the treatment of these tumors is warranted by targeting PNKP enzyme. PNKP functions both in NHEJ and single-strand break repair pathways (Dumitrache and McKinnon, 2017) and altering both could support the synthetic growth defect observed. Since PNKP inhibition is already exploited to kill other cancers (Mereniuk et al., 2013), we suggest that it be used to sensitize pHGG cells to current radio/chemotherapeutic regimens (Pinto et al., 2021), for which there is very limited response. Moreover, the specificity towards H3.3 K27M and G34R mutant cells may limit toxicity-related side effects in future treatments and opens up the possibility of employing a similar strategy in other cancers bearing the same H3.3 mutations (Deshmukh et al., 2021).

## Experimental procedures

### Human cell lines

U2OS (human osteosarcoma, female, American Type Culture Collection ATCC HTB-96) and HeLa cells (human cervical carcinoma, female, ATCC CCL-2) were cultured in Dulbecco’s modified Eagle’s medium DMEM Gluta-Max (Life Technologies) supplemented with 10% fetal bovine serum (Eurobio) and antibiotics (100 U/ml penicillin, 100 μg/ml streptomycin, Life Technologies) and maintained at 37 °C under 5% CO_2_ in a humified incubator. U2OS cells stably expressing SNAP-tagged wild-type or mutant H3.3, and U2OS LacO H3.3-SNAP cells with integrated 256 tandem LacO repeats and stably expressing SNAP-tagged wild-type H3.3(Adam et al., 2016) were cultured in the same medium supplemented with 100 μg/ml G418 (Life Technologies). Flp-In T-REx HEK293 cells (ThermoFisher) were cultured in DMEM (Gibco), supplemented with 10% FBS (Wisent) and 1% penstrep (Invitrogen), at 37°C in 5% CO_2_.

### Generation of U2OS stable cell lines

U2OS cells stably expressing C-terminal, SNAP-tagged H3.3, either wild-type, K27M, G34R, G34V, G34W or K36M mutated were generated by transfection of plasmid encoding wildtype or mutated H3.3 and selection of clones in limiting dilution in medium supplemented with G418 (Life Technologies) starting 48 hours after transfection. To verify the presence of mutations in the clones, genomic DNA was extracted and subjected to PCR amplification with the following primers: 5’-TGGCAGTACATCTACGTATTAGTCA-3’ (upstream of the CMV promoter) and 5’-GCTGGTGAAAGTAGGCGTTG-3’ (N-terminal to SNAP). The amplification product was verified by Sanger sequencing (GATC Biotech). Single clones harboring each H3.3 mutation were expanded and evaluated for levels of expression of the exogenous SNAP-tagged H3.3 proteins and for the presence of histone PTM alterations described in tumor samples.

### Primary pediatric human glioma cell lines

SF9402 and SF9427 (wild-type H3.3) cell lines were provided by C. D. James at Northwestern University (Chicago, USA) and cultured as previously reported (Hashizume et al., 2014). SU-DIPG-XVII (H3.3K27M) was provided by P. Knoepfler (University of California, Davis, USA) (Chen et al., 2020b). HSJD-002-GBM cells (H3.3G34R) were obtained from A. Carcaboso (Institut de Recerca Sant Joan de Deu, Barcelona, Spain). HSJD-002-GBM were cultured in Tumor Stem Medium (Nagaraja et al., 2017), which contains DMEM/F12 1:1 (Invitrogen), Neurobasal-A (Invitrogen), 10 mM HEPES (Invitrogen), 1× MEM sodium pyruvate (Invitrogen), 1× MEM non-essential amino acids (Invitrogen), 1% GlutaMax (Invitrogen), 20 ng/mL human basic fibroblast growth factor (CliniSciences), 20 ng/mL human epidermal growth factor (CliniSciences), 20 ng/mL human platelet-derived growth factor (PDGF)-A and PDGF-B (CliniSciences), 10 ng/mL heparin (StemCell Technologies), and 1x B27 without Vitamin A (Invitrogen). DIPG lines were generally grown in suspension flasks as tumorspheres, except when they underwent transfection and proliferation assay, for which they were dissociated and plated on plates coated with laminin (10 μg/mL, Sigma-Aldrich). MGBM1 cells (H3.3G34R) were obtained from N. Jabado (McGill University, Montreal, Canada) (Bender et al., 2013) and cultured in DMEM Gluta-Max supplemented with 10% FBS and antibiotics (100 U/ml penicillin, 100 μg/ml streptomycin). All glioma cells were maintained at 37 °C under 5% CO_2_ in a humified incubator and verified for expression of the expected H3.3 proteins by Western blot analysis with antibodies raised against H3.3K27M or G34R (see Antibody list for details).

### Drug treatments and inhibitors

Camptothecin (CPT, Sigma-Aldrich) was used at 0.1 μM for 3 h, or at 1 μM for 1 or 3 h for iPOND in human cells, and at 5 or 10 μM in yeast cells; hydroxyurea (HU, Sigma-Aldrich) at 2 mM for 3 h; mitomycin C (MMC, Sigma-Aldrich) at 200 ng/mL and 25 ng/mL for 24 h for repair foci analyses and metaphase spreads, respectively; bleomycin (Bleo, Sigma-Aldrich) at 20 μg/mL for 3 h. An overnight treatment with 2 mM Thymidine followed by 3 h release in fresh medium was used to enrich cells in S phase for iPOND experiments (70-75% of cells were in S phase as evaluated by FACS). The EZH2 inhibitor GSK126 (EZH2i, Selleckchem) was used at 1 μM for 72 h.

### Plasmids and site directed mutagenesis

The H3F3A and H3F3B human cDNA sequences (GenScript) were cloned by using ClaI and EcoRI restriction enzymes into the pSNAPm plasmid (New England Biolabs), with the SNAP tag in the C-terminus of the insert. These plasmids were subjected to directed mutagenesis to introduce the cancer-associated mutations (Behjati et al., 2013; Schwartzentruber et al., 2012; Wu et al., 2012) (see Table 2 for details of the mutations, primers used and genes involved). Generation of the mutated plasmids was verified by Sanger sequencing (GATC Biotech).

**Table 1.**
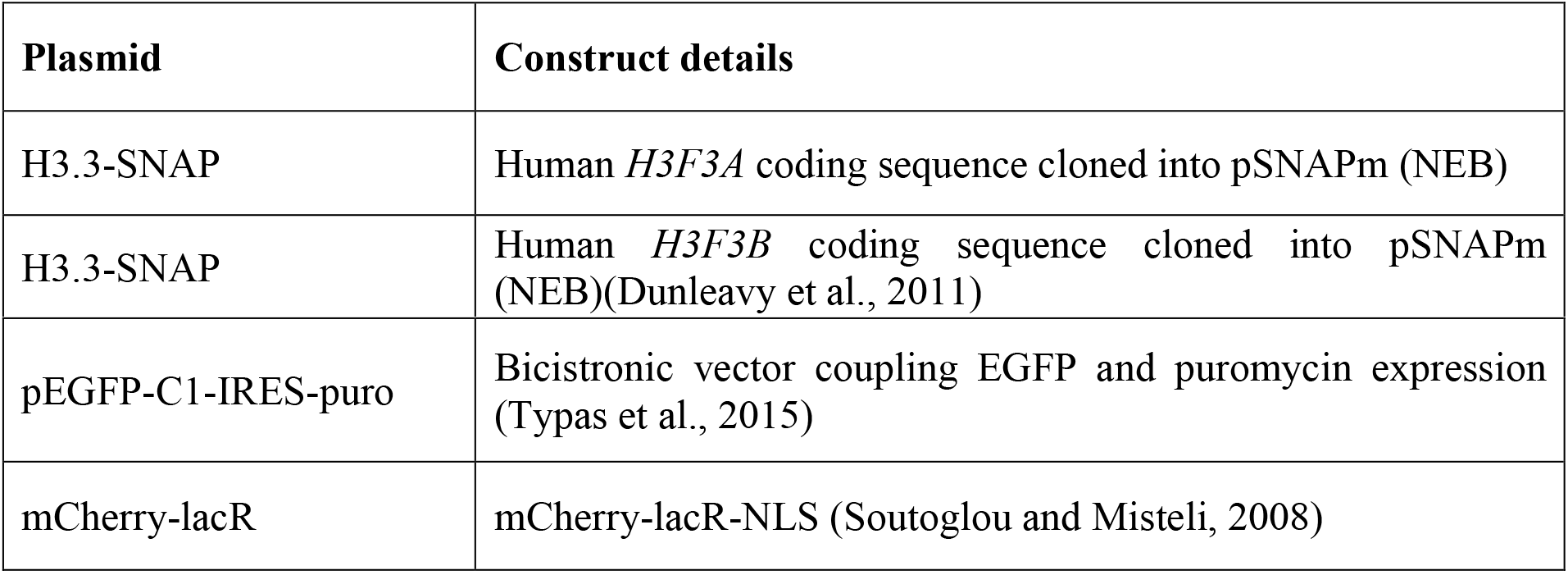
Plasmids used in this study.

**Table 2:**
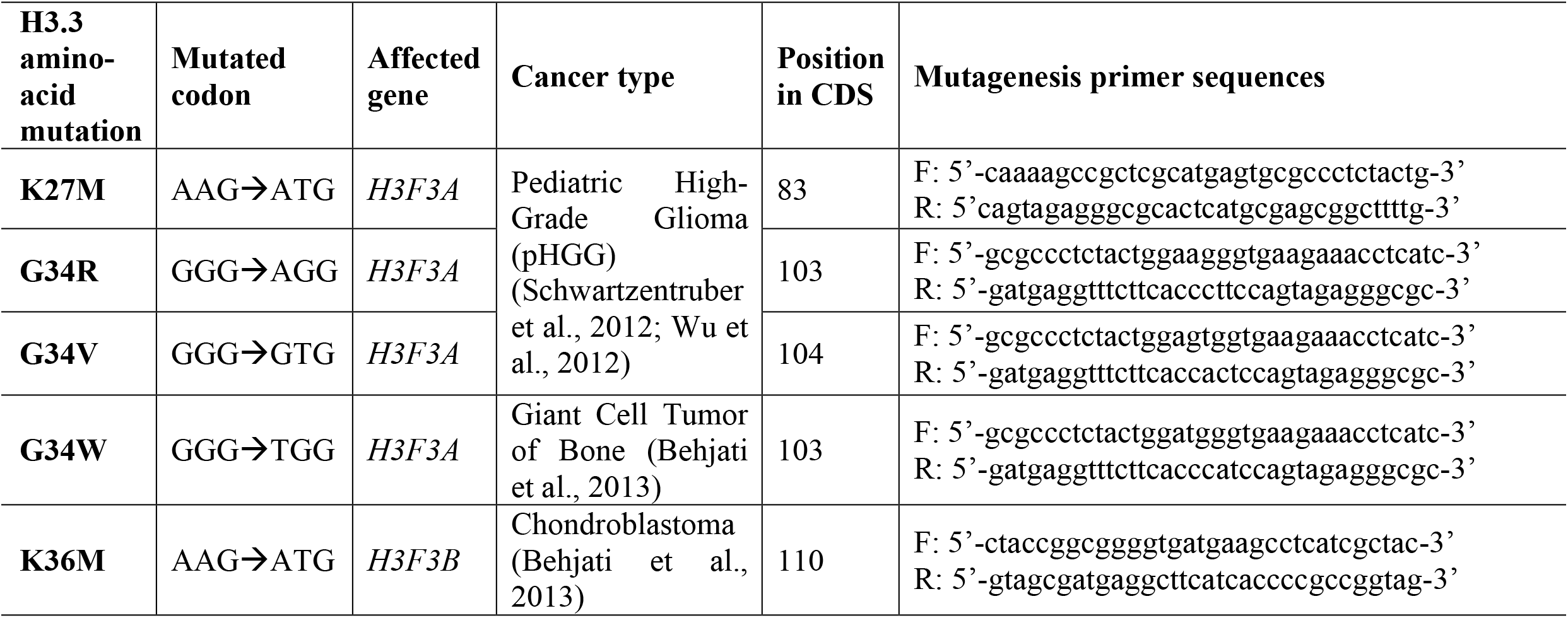
Point mutations in H3.3 coding genes: primers used and associated cancers. F = Forward, R = Reverse, CDS = Coding DNA Sequence.

### Immunofluorescence, image acquisition and analysis

Cells grown on glass coverslips (VWR) were either fixed directly with 2% paraformaldehyde (PFA) and permeabilized with 0.2% Triton X-100 in PBS or pre-extracted before fixation with 0.5% Triton X-100 in CSK buffer (Cytoskeletal buffer: 10 mM PIPES pH 7.0, 100 mM NaCl, 300 mM sucrose, 3 mM MgCl2) to remove soluble proteins (not bound to chromatin) and then fixed with 2% PFA. Samples were blocked in 5% Bovine Serum Albumin (BSA, Sigma-Aldrich) in PBS supplemented with 0.1% Tween 20 (Euromedex) before incubation with primary antibodies and secondary antibodies conjugated to Alexa Fluor 488 or 568 (Invitrogen). Coverslips were mounted in Vectashield medium with DAPI (Vector Laboratories) and observed with a Leica DMI6000 epifluorescence microscope using a Plan-Apochromat 40x/1.3 or 63x/1.4 oil objective. Images were captured using a CCD camera (Photometrics) and Metamorph software. Images were mounted with Adobe Photoshop applying the same treatment of fluorescence levels to all images from the same experiment. Fiji software was used for image analyses using custom macros. Nuclei were delineated based on DAPI staining, and CPT-damaged RFs based on gH2A.X staining. S phase, replicating cells were discriminated based on EdU staining. The position of the LacO array was determined based on mCherry-LacR signal. DNA repair and PLA foci were identified and counted by using the find maxima function (Fiji software), on maximum intensity z-projections in the case of PLA foci. At least 70 cells/sample were scored in each experiment. Results of automatic foci counting were graphed as number of foci per cell or as number of cells with more than 5 or 10 DNA repair foci that was set as a threshold.

### Ethynyl-deoxyUridine (EdU) labeling of S phase cells

For discrimination of S phase cells, 10 μM 5-Ethynyl-2’-deoxyUridine (EdU, Sigma-Aldrich) was incorporated into cells for 15 minutes prior to DNA damage treatment and fixation. EdU was revealed using Click-It EdU Imaging kit (Invitrogen) according to manufacturer’s instructions.

### Random plasmid integration assay

The assay was performed as in (Galanty et al., 2009). Briefly, cells grown in 6-well plates were transfected with siRNAs and, later the same day, the cells were transfected with 2 μg/well gel-purified FspI-BspDI-linearized pEGFP-C1-IRES-puro plasmid (provided by van Attikum lab, Leiden, the Netherlands). The cells were transfected once more with siRNAs the following day. Cells were collected 48 h later, counted and seeded in 10 cm diameter dishes either lacking or containing 0.375 μg/mL puromycin. The transfection efficiency was determined on the same day by FACS analysis of EGFP-positive cells. The cell dishes were incubated at 37°C to allow colony formation and medium was refreshed on day 4 and 8. On day 10-12, the cells were stained with 0.5 % Crystal Violet (Sigma-Aldrich)/20% ethanol solution to score colonies with more than 50 cells. Random plasmid integration events on the puromycin-containing plates were normalized to the plating efficiency (plate without puromycin) and to the transfection efficiency.

### Fission yeast strains and genetic analyses

*Schizosaccharomyces pombe* strains (Table 3) containing point mutations in histone H3, K27M in *hht2*^+^, G34R and G34V in *hht3*^+^, were generated by a PCR-based module method. *pnk1Δ* and *xrc4*Δ strains were derived from the fission yeast deletion library and the gene deletions were verified by PCR. All other strains were constructed through genetic crosses. For serial dilution plating assays (spot assays), ten-fold dilutions of a mid-log phase culture were plated on the indicated medium and grown for 3 days at 30°C. For proliferation assays, overnight liquid *S. pombe* cultures were grown to saturation in YES media (Yeast Extract with Supplement). Saturated cultures were equilibrated to an OD_600_ of 1.0, arrayed in a 96-well microtiter plate, and pinned in quadruplicate to achieve a 384-colony density (i.e. 4 technical replicates for each position of the microtiter plate) using a Singer RoToR robot (Singer Instruments, Inc. Somerset UK). Strains were grown on YES solid agar media with the indicated camptothecin concentrations. Plates with pinned colonies were incubated at 30°C and scanned every 96 minutes for growth curves. Colony density was measured using screenmill (Bryant et al., 2019)_and the values of the 4 technical replicates were averaged. Care was taken to minimize systematic bias in experiments (e.g. by distributing strains evenly throughout the 96-well plate to minimize position and neighboring strain effects).

**Table 3:** Fission yeast strains used in this study.

|  |  |
| --- | --- |
| SPJ1577 | *h<sup>+</sup> leu1 ura4 ade6* |
| SPJ4742 | *h<sup>+</sup> leu1 ura4 ade6 hht3-G34R-Flag::hphMX6* |
| SPJ4738 | *h<sup>+</sup> leu1 ura4 ade6 hht3-G34V-Flag::hphMX6* |
| SPJ5245 | *h<sup>+</sup> leu1 ura4 ade6 hht2-K27M::NatMX6* |
| CS2653 | *h<sup>+</sup> leu1 ura4 ade6 xrc4Δ::kanMX6* |
| CS2656 | *h<sup>+</sup> leu1 ura4 ade6 hht3-G34R-Flag::hphMX6 xrc4Δ::kanMX6* |
| CS2655 | *h<sup>+</sup> leu1 ura4 ade6 hht3-G34V-Flag::hphMX6 xrc4Δ::kanMX6* |
| SPJ5276 | *h<sup>+</sup> leu1 ura4 ade6 hht2-K27M::NatMX6 xrc4Δ::kanMX6* |
| CS2669 | *h<sup>+</sup> leu1 ura4 ade6 hht3-G34R-Flag::hphMX6 pnk1Δ::kanMX6* |
| SPJ5454 | *h<sup>+</sup> leu1 ura4 ade6 hht3-G34R-Flag::hphMX6 xrc4Δ::kanMX6 pnk1Δ::kanMX6* |

**Table 3:** Antibodies used. HRP: HorseRadish Peroxidase; IF: Immunofluorescence; PLA: Proximity Ligation assay WB: western blot

|  | **Antibody target** | **Species** | **Supplier (Reference)** | **Dilution** | **Application** |
| --- | --- | --- | --- | --- | --- |
| Primary antibodies | 53BP1 | Rabbit | Novus Biologicals (NB100-304) | 1:1000 | WB<br>IF |
|  | Biotin | Rabbit | Bethyl Laboratories (BETA150-109A) | 1:500 | PLA |
|  | CAF-1 p60 | Rabbit | Abcam (ab109442) | 1:1000 | WB |
|  | FANCD2 | Rabbit | Novus Biologicals (NB100-182) | 1:500 | WB<br>IF |
| | γH2A.X | Rabbit | Cell Signaling Technology (2577) | 1:1000 | WB |
| | γH2A.X | Mouse | EMD Millipore (05-636) | 1:1000 | IF<br>PLA |
|  | H3 | Rabbit | Abcam (ab1791) | 1:1000 | WB |
|  | H3.3 | Rabbit | EMD Millipore (09-838) | 1:1000 | WB |
|  | H3.3K27M | Rabbit | EMD Millipore (ABE419) | 1:1000 | WB |
|  | H3.3G34R | Rabbit | Cambridge Research Biochemicals (crb2005185f) | 1:250 | WB |
|  | H3.3G34V | Rabbit | Cambridge Research Biochemicals (crb2005186f) | 1:1000 | WB |
|  | H3.3K36M | Rabbit | RevMab (31-1085-00) | 1:1000 | WB |
|  | H3K27me3 | Rabbit | Cell Signaling Technology (9733) | 1 :250 | WB |
|  | H3K36me3 | Rabbit | Abcam (ab9050) | 1:2000 | WB |
|  | H4 | Rabbit | Actif Motif (61300) | 1:1000 | WB |
|  | H4K20me2 | Rabbit | Abcam (ab9052) | 1:1000<br>1:1000 | WB<br>IF |
|  | PCNA | Mouse | Dako (M0879) | 1:000 | WB |
|  | PNKP | Rabbit | Sigma-Aldrich (HPA006782) | 1:500 | WB |
|  | RAD51 | Rabbit | Santa Cruz Biotechnology sc-8349 (Clone H-92) | 1:500<br>1:250 | WB<br>IF |
|  | RNF168 | Rabbit | EMD Millipore (ABE367) | 1:1000 | WB |
|  | SETD2 | Rabbit | Abcam (ab31358) | 1:250 | WB |
|  | SNAP | Rabbit | Pierce Antibodies (CAB4255) | 1:500<br>1:1000 | WB<br>IF |
|  | Tubulin | Mouse | Sigma-Aldrich (T9026) | 1:10000 | WB |
|  | XRCC4 | Rabbit | Sigma-Aldrich (HPA006801) | 1:500 | WB |
| Secondary antibodies | Mouse Alexa Fluor 488 | Goat | Invitrogen (A11029) | 1:1000 | IF |
|  | Mouse Alexa Fluor 568 | Goat | Invitrogen (A11031) | 1:1000 | IF |
|  | Rabbit Alexa Fluor 488 | Goat | Invitrogen (A11034) | 1:1000 | IF |
|  | Rabbit Alexa Fluor 568 | Goat | Invitrogen (A11036) | 1:1000 | IF |
|  | Mouse HRP | Goat | Jackson Laboratories 115-035-068 | 1:10000 | WB |
|  | Rabbit HRP | Donkey | Jackson Laboratories 711-035-152 | 1:10000 | WB |

### Metaphase spreads

To prepare metaphase spreads, Colcemid (Gibco) was added to the culture medium at 0.1 μg/ml for 3 h before collecting the cells. Cells were washed in PBS, trypsinized, centrifuged at 1200 rpm for 5 min and resuspended in 75 mM KCl for 15 min at 37°C. Cells were then fixed with fresh ethanol/acetic acid (v:v=3:1) at −20°C overnight. The next day, cells were centrifuged at 1200 rpm for 5 min and resuspended in fresh fixative before dropping onto slides and then air dried. Slides were mounted with VECTASHIELD Antifade Mounting Medium containing DAPI (Vector laboratories) and metaphase spreads were examined using an Axio Observer Z1 epifluorescence microscope equipped with an ORCA-ER camera (Hamamatsu). Mitomycin (MMC) was added at a concentration of 25 ng/mL for 48 hours before harvesting cells for metaphase spread preparation. At least 30 metaphase spreads were scored per sample in each experiment for the presence of radial chromosomes.

### Mutational signature analysis on primary pHGG samples

pHGG samples for single nucleotide variant (SNV) mutational signature analysis were acquired from previously published data available under EGAS00001000575, EGAS00001001139, EGAS00001000572 and EGAS00001000192. Novel data was generated from samples obtained from the DIPG-BATs clinical trial (NCT01182350), the Dana-Farber Tissue Bank or collaborating institutions, under protocols approved by the institutional review board of the Dana-Farber/Harvard Cancer Center with informed consent (DFCI protocols 10417, 10201 and DFCI 19293). DNA was extracted from single Diffuse Midline Glioma cores, pHGG biopsies and autopsy samples using Qiagen AllPrep DNA/RNA extraction kits. For whole-genome sequencing, genomic DNA was fragmented and prepared for sequencing to 60X depth on an Illumina HiSeq 2000 instrument. Reads from both novel and published data were aligned to the reference genome hg19/ GRCh37 with BWA83, duplicate-marked, and indexed using SAMtools and Picard. Base quality score was bias adjusted for flowcell, lane, dinucleotide context, and machine cycle and recalibrated, and local realignment around insertions or deletions (indels) was achieved using the Genome Analysis Toolkit. SNV signature analysis was performed using Palimpsest (https://github.com/FunGeST/Palimpsest) on a VCF containing somatic mutations identified by Mutect2.

### Extraction of cellular proteins and Western blot analysis

Total extracts were obtained by scraping cells in Laemmli buffer (50 mM Tris HCl pH 6.8, 1.6% Sodium Dodecyl Sulfate (SDS), 8% glycerol, 4%β-mercaptoethanol, 0.0025% bromophenol blue) followed by 5-10 min denaturation at 95 °C.

For western blot analysis, extracts along with molecular weight markers (Precision plus protein Kaleidoscope standards, Bio-Rad) were run on 4%–20% Mini-PROTEAN TGX gels (Bio-Rad) in running buffer (200 mM glycine, 25 mM Tris, 0.1% SDS) and transferred onto nitrocellulose membranes (Protran) with a Trans-Blot SD semi-dry or wet transfer cell (Bio-Rad). Proteins of interest were probed using the appropriate primary and Horse Radish Peroxidase (HRP)-conjugated secondary antibodies (Jackson Immunoresearch), detected using SuperSignal West Pico or Femto chemiluminescence substrates (Pierce). The resulting signal was visualized on hyperfilms MP (Amersham) with a film processor (SRX105, Konica). ImageJ software was used for densitometric quantification of protein bands.

### siRNA and plasmid transfections

siRNAs purchased from Eurofins MWG Operon or Sigma-Aldrich (Table 4) were transfected into cells using Lipofectamine RNAiMAX (Invitrogen) following manufacturer’s instructions. Cells were analyzed and/or harvested 48 to 72 h post-transfection except for proliferation assays, where cells were analyzed over a 7-day period after transfection.

**Table 4:**
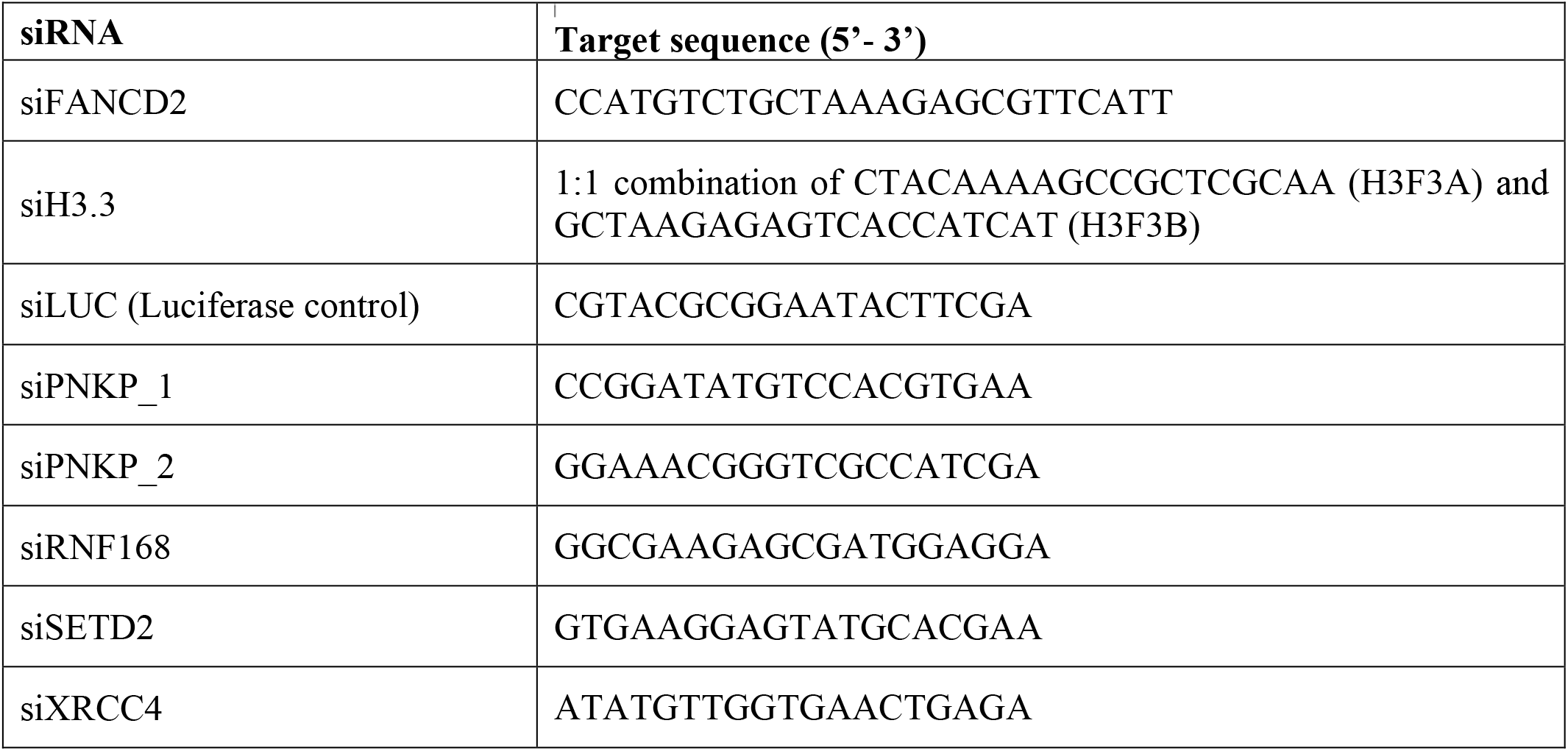
siRNA sequences.

Cells were transfected with plasmid DNA (see plasmid section) using Lipofectamine 2000 (Invitrogen) according to manufacturer’s instructions.

### SNAP labeling of newly synthesized histones

For labeling newly synthesized SNAP-tagged histones (Bodor et al., 2012), parental histones were quenched with 10 μM SNAP-cell Block (NEB) for 30 minutes in culture medium followed by 30-min wash in fresh medium and a 2-h chase. To mark S phase cells/replication forks, EdU was incorporated for 30 minutes at the end of the chase period. The new SNAP-tagged histones synthesized during the chase were fluorescently labelled with 4 μM of the green-fluorescent reagent SNAP-cell Oregon green (New England Biolabs) during a 15-min pulse step followed by 30-min wash in fresh medium. Alternatively, when combined with Proximity Ligation Assay (PLA), new SNAP-tagged histones were pulse-labeled for 30 min with 5 μM final SNAP-biotin (New England Biolabs) diluted 1:200 in 10% Duolink blocking buffer (Sigma-Aldrich) in PBS. After washings, soluble proteins were removed by permeabilization with 0.5% Triton X-100 in cytoskeleton (CSK) buffer, and cells were fixed and processed for immunostaining or PLA.

### Proximity Ligation assay (PLA)

PLA (Söderberg et al., 2006) was performed to detect colocalization foci between newly synthesized H3.3-SNAP and gH2A.X at camptothecin-damaged replication forks. The Duolink^®^ In Situ PLA^®^ detection kit (Sigma) was used following manufacturer’s recommendations. Briefly, cells on glass coverslips (VWR) were incubated 1 h at 37 °C in Duolink blocking buffer (Sigma-Aldrich) and then for 1 h at room temperature with a mix of the two primary antibodies directed against the target proteins (anti-biotin to detect new H3.3-SNAP-biotin and anti-gH2A.X to detect sites of DNA damage) diluted in antibody dilution reagent (Sigma-Aldrich). Coverslips were then incubated for 1 h at 37 °C with secondary antibodies each harboring a PLA probe (Duolink In Situ PLA MINUS/PLUS probes, Sigma-Aldrich). The PLA probes that bind to the constant regions of the primary antibodies contain a unique DNA strand. If the proteins of interest interact with each other, the DNA probes hybridize to make circular DNA during the 30 min ligation step at 37 °C. The resulting circle DNA can be amplified (1 h 40 min amplification at 37 °C, Duolink In Situ Detection Reagents Green, Sigma-Aldrich) and visualized by fluorescently labeled complementary oligonucleotide probes incorporation. Coverslips were mounted in Duolink In Situ Mounting Medium with DAPI (Sigma-Aldrich). To study PLA foci in S phase cells, EdU labeling by click chemistry was performed before the blocking step.

### Isolation of proteins on nascent DNA (iPOND)

iPOND was performed largely as described previously (Sirbu et al., 2012), with the following modifications. A total of 3 × 10^7^ logarithmically growing cells per sample were labeled with 10 μM EdU for 15 min. Following EdU incorporation, cells were treated or not with camptothecin, fixed with 1% formaldehyde for 15 min at room temperature, followed by 5-min incubation with 0.125 M glycine to quench the formaldehyde. Cells were harvested by scraping, washed three times with PBS, flash frozen in liquid nitrogen and kept at −80 °C. Within two weeks, samples were processed for EdU-based pulldown and purification of replication fork-associated proteins. Briefly, click chemistry reactions were performed on pre-permeabilized samples to conjugate biotin to the EdU-labeled DNA by using Biotin Picolyl azide (Sigma Aldrich). Sonication was performed with a Bioruptor Pico sonicator (Diagenode) and DNA shearing was evaluated on an agarose gel. Shearing for optimal detection of the proteins of interest was set to an average DNA fragment size of 800 bp. Total input samples were taken after sonication and clearing of samples and kept at −20 °C until loading on SDS-PAGE gels. Streptavidin beads (Dynabeads MyOne Streptavidin-C1, Life technologies) were used to capture the biotin-conjugated DNA-protein complexes. Captured complexes were washed extensively using SDS and high-salt wash buffers. Purified replication fork proteins were eluted under reducing conditions by boiling in Laemmli sample buffer for 5 min. Total input and capture samples corresponding to equal amounts of cells were resolved on SDS-PAGE gels and analyzed by western blot.

### Flow cytometry and cell cycle analysis

Cells were fixed in ice-cold 70% ethanol before DNA staining with 50 μg/mL propidium iodide (Sigma-Aldrich) in PBS containing 0.05% Tween 20 and 0.5 mg/mL RNase A (USB/Affymetrix). DNA content was analyzed by flow cytometry using a FACSCalibur Flow Cytometer (BD Biosciences) and FlowJo Software (TreeStar).

### Proximity biotinylation

Flp-In T-REx HEK293 cells were induced for 24 h with 1 μg/ml doxycycline and 50 μM biotin and snap frozen before lysis in 10 ml lysis buffer (50 mM Tris-HCl pH 7.5, 150 mM NaCl, 1 mM EDTA, 1 mM EGTA, 1% Triton X-100, 0.1% SDS, 1:500 protease inhibitor cocktail (Sigma-Aldrich), 1:1,000 benzonase nuclease (Novagen)). Samples were rotated at 4°C for 1 h, briefly sonicated and centrifuged at 45,000 x g for 30 min at 4°C. Cleared supernatants were added to 30 μl of packed, pre-equilibrated Streptavidin sepharose beads (GE Healthcare) and incubated for 3 hours at 4°C with end-over-end rotation. Beads were centrifuged at 2,000 rpm for 2 min, and in a fresh tube washed twice with 1 mL of lysis buffer and twice with 1 mL of 50 mM ammonium bicarbonate pH 8.3. Beads were transferred in ammonium bicarbonate to a fresh centrifuge tube and washed two more times. On-bead tryptic digestion was performed with 1 μg MS-grade TPCK trypsin (Promega, Madison, WI) dissolved in 200 μl of 50 mM ammonium bicarbonate pH 8.3 overnight at 37°C. The following morning, 0.5 μg MS-grade TPCK trypsin was added, and beads were incubated 2 additional hours at 37°C. Beads were pelleted by centrifugation at 2,000 x g for 2 min, and the supernatant was transferred to a fresh Eppendorf tube. Beads were washed twice with 150 μl of 50 mM ammonium bicarbonate, and the washes were pooled with the first eluate. The sample was lyophilized and resuspended in buffer A (0.1% formic acid). 1/5th of the sample was analyzed per mass spectrometry run.

Mass spectrometry was performed and analyzed as previously described (Gupta et al., 2015). Briefly, high performance liquid chromatography was conducted using a 2 cm pre-column (Acclaim PepMap 50 mm x 100 μm inner diameter (ID)), and 50 cm analytical column (Acclaim PepMap, 500 mm x 75 μm diameter; C18; 2 μm; 100 Å, Thermo Fisher Scientific, Waltham, MA), running a 120 min reversed-phase buffer gradient at 225 nl/min on a Proxeon EASY-nLC 1000 pump in-line with a Thermo Q-Exactive HF quadrupole-Orbitrap mass spectrometer. A parent ion scan was performed using a resolving power of 60,000, then up to the twenty most intense peaks were selected for MS/MS (minimum ion count of 1,000 for activation) using higher energy collision induced dissociation (HCD) fragmentation. Dynamic exclusion was activated such that MS/MS of the same m/z (within a range of 10 ppm; exclusion list size = 500) detected twice within 5 sec were excluded from analysis for 15 sec. For protein identification, Thermo .RAW files were converted to the .mzXML format using Proteowizard (Kessner et al., 2008), then searched using X!Tandem (Craig and Beavis, 2004) and COMET (Eng et al., 2013) against the human Human RefSeq Version 45 database (containing 36,113 entries). Data were analyzed using the trans-proteomic pipeline (TPP) (Deutsch et al., 2010; Pedrioli, 2010) via the ProHits software suite (v3.3) (Liu et al., 2010). Search parameters specified a parent ion mass tolerance of 10 ppm, and an MS/MS fragment ion tolerance of 0.4 Da, with up to 2 missed cleavages allowed for trypsin. Variable modifications of + 16@M and W, +32@M and W, +42@N-terminus, and +1@N and Q were allowed. Proteins identified with an iProphet cut-off of 0.9 (corresponding to ≤1% FDR) and at least two unique peptides were analyzed with SAINT Express v.3.3.1. Ten control runs (from cells expressing the FlagBirA* epitope tag) were collapsed to the two highest spectral counts for each prey and compared to the two biological and two technical replicates of histone BioID. High confidence interactors were defined as those with Bayesian false discovery rate (BFDR) ≤0.01.

### Human cell proliferation assays

The effect of PNKP knockdown on cell proliferation in human cells was measured as follows: 24 h after siRNA transfection, cells were seeded in 60-mm diameter tissue culture plates (20 000 cells/plate for U2OS, 40 000 to 80 000 for pHGG cells). pHGG cell lines were transfected with siRNAs twice, both 48 h and 24 h before seeding. Cell viability was assessed after 3, 5 and 7 days in culture by staining with trypan blue (Invitrogen) or counting with an automated cell counter for viable cells (Fluidlab R-300 cell counter, Anvajo).

### Statistical analysis

Statistical analyses were carried out using Graphpad Prism software. P values for mean comparisons between two groups were calculated with a Student’s t-test with Welch’s correction when necessary. Multiple comparisons were performed by one- or two-way ANOVA with Bonferroni, Tukey’s or Dunnett’s post-tests or using the non-parametric Kruskal-Wallis test in case of non-gaussian distributions. Comparisons of proliferation curves were based on non-linear regression with a polynomial quadratic model. ns: non-significant, *p < 0.05, **p < 0.01, ***p < 0.001, ****: p<0.0001. Statistical parameters including sample size (n) and dispersion of the data (SD or SEM) are indicated in the figure legends.

### Resource and data availability

Raw mass-spectrometry data is available for download from https://massive.ucsd.edu/, accession numbers MSV000087736 (reviewer login: username, MSV000087736_reviewer; password, HISTH3_BioID) and MSV000089173 (reviewer login: username, MSV000089173_reviewer; password: H3_mutants). Whole genome sequencing data is available under EGAS00001000575, EGAS00001001139, EGAS00001000572, EGAS00001000192, and under dbGaP accession number phs002380.v1.p1 (Dubois et al., 2022). All other data are available in the main text or the supplemental information.

## Acknowledgments

We thank members of our laboratory, M. Touat and F. Bielle for stimulating discussions, S. Ait-Si-Ali and P-A. Defossez for critical reading of the manuscript, A. Carcaboso, N. Jabado, C.D. James, P. Knoepfler and H. van Attikum for sharing pHGG cell lines and reagents. We acknowledge the imaging platform of the Epigenetics and Cell Fate Centre. Research in R.R. lab was supported by the National Institutes of Health (R35GM118180). Research in S.J. lab was supported by the National Institutes of Health (R35GM126910). Work in P.B. lab was funded by Alex’s Lemonade Stand Foundation and Prayers from Maria. Work in R.B. lab was funded by the Gray Matters Brain Cancer Foundation and the Brown Fund for Innovation in Cancer Informatics. Work in V.N. lab was supported by the European Research Council (ERC-2014-StG-638898). Work in C.H. lab was supported by the Canadian Cancer Society Research Institute (702296 and 706160) and the Canadian Institutes of Health Research (PJT-156407). Research in S.E.P. lab was supported by the European Research Council (ERC-2013-StG-336427 and ERC-2018-CoG-818625), the French National Research Agency (ANR-12-JSV6-0002-01 and ANR-18-CE12-0017-01), and the Labex “Who am I?” (ANR-11-LABX-0071, ANR-18-IDEX-0001). B.R. was a Marie Curie fellow (H2020-MSCA-IF-2017-799101). G.G. is supported by a PhD grant from the Fondation pour la Recherche Medicale (ECO202106013684). S.E.P. is an EMBO Young Investigator.

## Author contributions

B.R., S.P., G.G. and O.C. performed most of the experiments. J.D. and S-K.B. generated U2OS cell lines expressing wild-type and mutant H3.3. B.K., R.S., B.R. and E.C. performed BioID experiments under the supervision of C.H. C-M.S., R.J.D.R. and T.T. performed fission yeast experiments with supervision from S.J and R.R. T.W. and V.B. supervised by V.N. performed metaphase spread analyses. A.C. and F.D. supervised by P.B. and R.B. performed mutational signature analyses. B.R. and S.E.P. conceived the study, supervised the work and wrote the manuscript. All authors edited and approved the final version of the manuscript.

