## Supplementary figures and legends for "Aberrant DNA repair is a vulnerability in histone H3.3-mutant brain tumors"

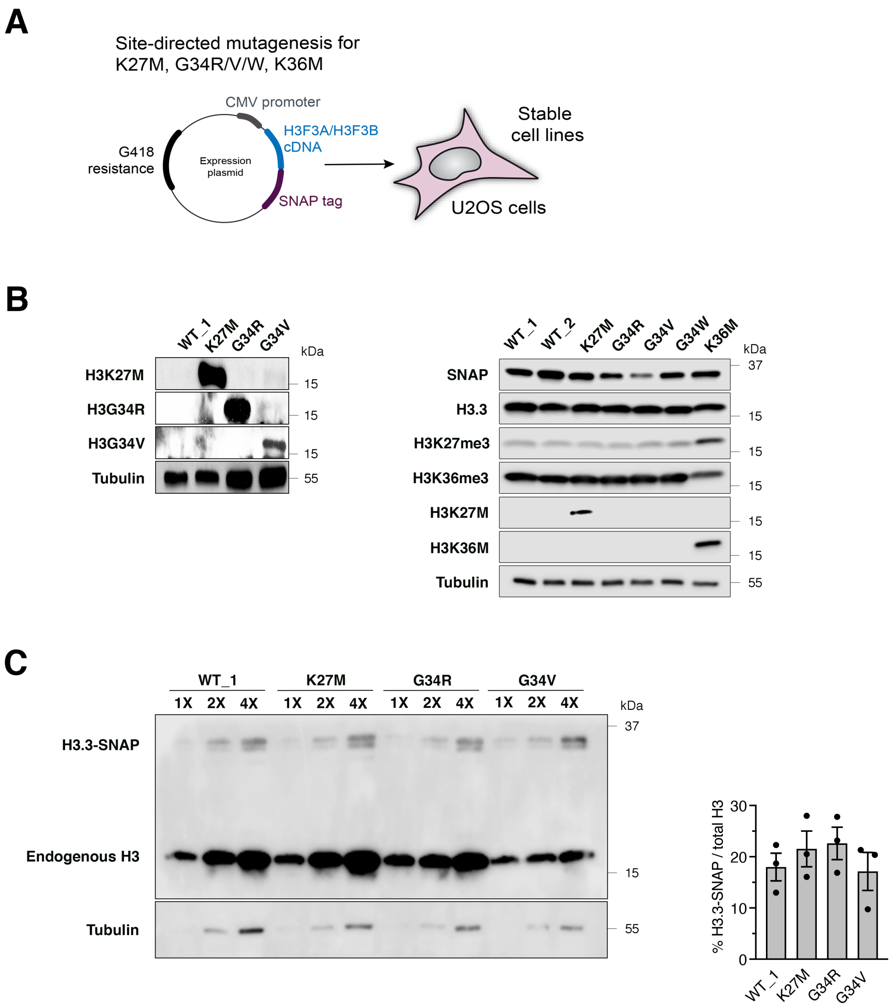


Supplemental Figure S1

**Figure S1. Characterization of stable U2OS cell lines expressing wild-type or mutant H3.3. Related to Figure 1.** (A) Graphical representation of the plasmid DNA used to express SNAP-tagged H3.3 proteins and to generate stable U2OS cell lines. (B) Western blot analysis of whole cell extracts from the generated cell lines showing expression of SNAP-tagged H3.3, the levels of H3K27me3 and H3K36me3 histone post-translational modifications and H3.3 mutations (with antibodies recognizing H3K27M, H3G34R, H3G34V, H3K36M). Tubulin is used as a loading control. The levels of H3K27me3 are not decreased in H3.3K27M as compared to wild-type H3.3 U2OS cells due to endogenous EZH2 inhibition, which significantly lowers the overall H3K27me3 levels in these cells. (C) Western blot analysis of SNAP-tagged H3.3 protein levels in whole cell extracts from the generated U2OS cell lines. The bar graph depicts the ratio of H3.3-SNAP to total histone H3 levels, which is expressed as a percentage. Mean ± SEM from three independent experiments.

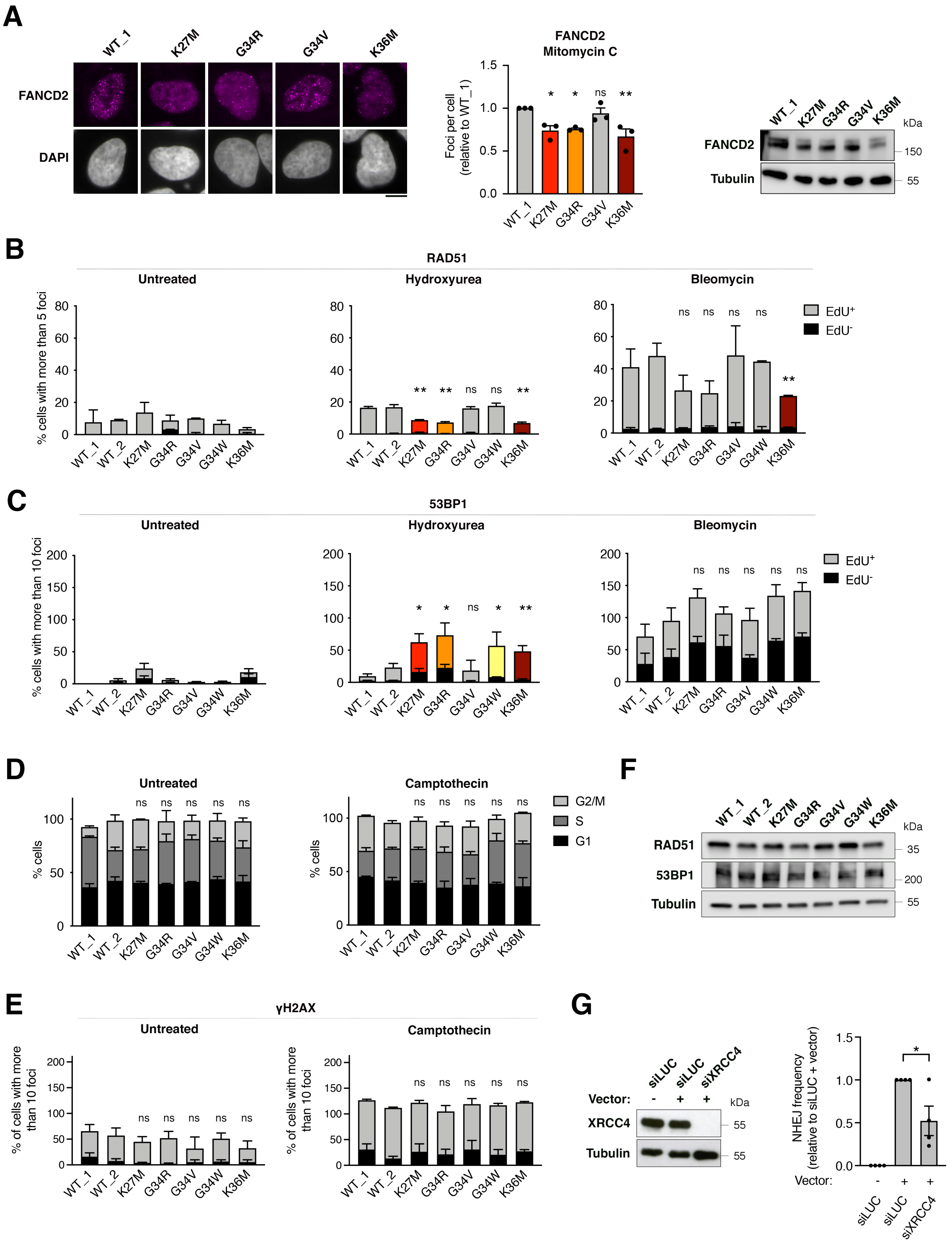


Supplemental Figure S2

**Figure S2. Analysis of the DNA repair defect in U2OS cells stably expressing H3.3 proteins. Related to Figure 1.** (A-C) Analysis of FANCD2 (A), RAD51 (B), 53BP1 (C) repair foci by immunofluorescence in U2OS cells stably expressing wild-type H3.3 (WT_1) or the indicated mutants and treated with mitomycin C (24 h, 200 ng/mL), hydroxyurea (3 h, 2 mM) or bleomycin (3 h, 20 µg/mL). Representative images of FANCD2 repair foci and FANCD2 protein levels in whole cell extracts are shown in (A) (Tubulin, loading control). Bar graphs depict the number of FANCD2 foci per cell (A), or the percentage of $EdU^+$ and $EdU^-$ cells harboring more than 5 (RAD51, B) or 10 (53BP1, C) foci. Mean ± SEM from three independent experiments. (D) Cell cycle distribution of U2OS cells stably expressing wild-type H3.3 (WT_1) or the indicated mutants untreated or treated with camptothecin (3 h, 0.1 µM). Bar graphs show the percentage of cells in G1, S and G2/M phases of the cell cycle with mean ± SEM from three independent experiments. (E) Analysis of γH2A.X damage foci by immunofluorescence in U2OS cells stably expressing wild-type H3.3 (WT_1) or the indicated mutants, untreated or treated with camptothecin (3 h, 0.1 µM). Mean ± SEM from three independent experiments, with n>134 per sample for each experiment. (F) Western blot analysis of whole cell extracts showing expression of RAD51, 53BP1 and Tubulin (loading control) in U2OS cells stably expressing wild-type H3.3 (WT_1) or the indicated mutants. (G) Analysis of NHEJ frequency by random plasmid integration assay in U2OS cells transfected with siRNA against Luciferase (siLUC, control) or against the core NHEJ factor XRCC4. Vector –, negative untransfected control. Statistical significance is calculated by one-way ANOVA (A and G), two-way ANOVA (B, C and E) or chi-square ($\chi^2$) test for distributions (D). *: $p< 0.05$; **: $p< 0.01$; ***: $p< 0.001$; ns: $p> 0.05$. Scale bars, 10 µm.

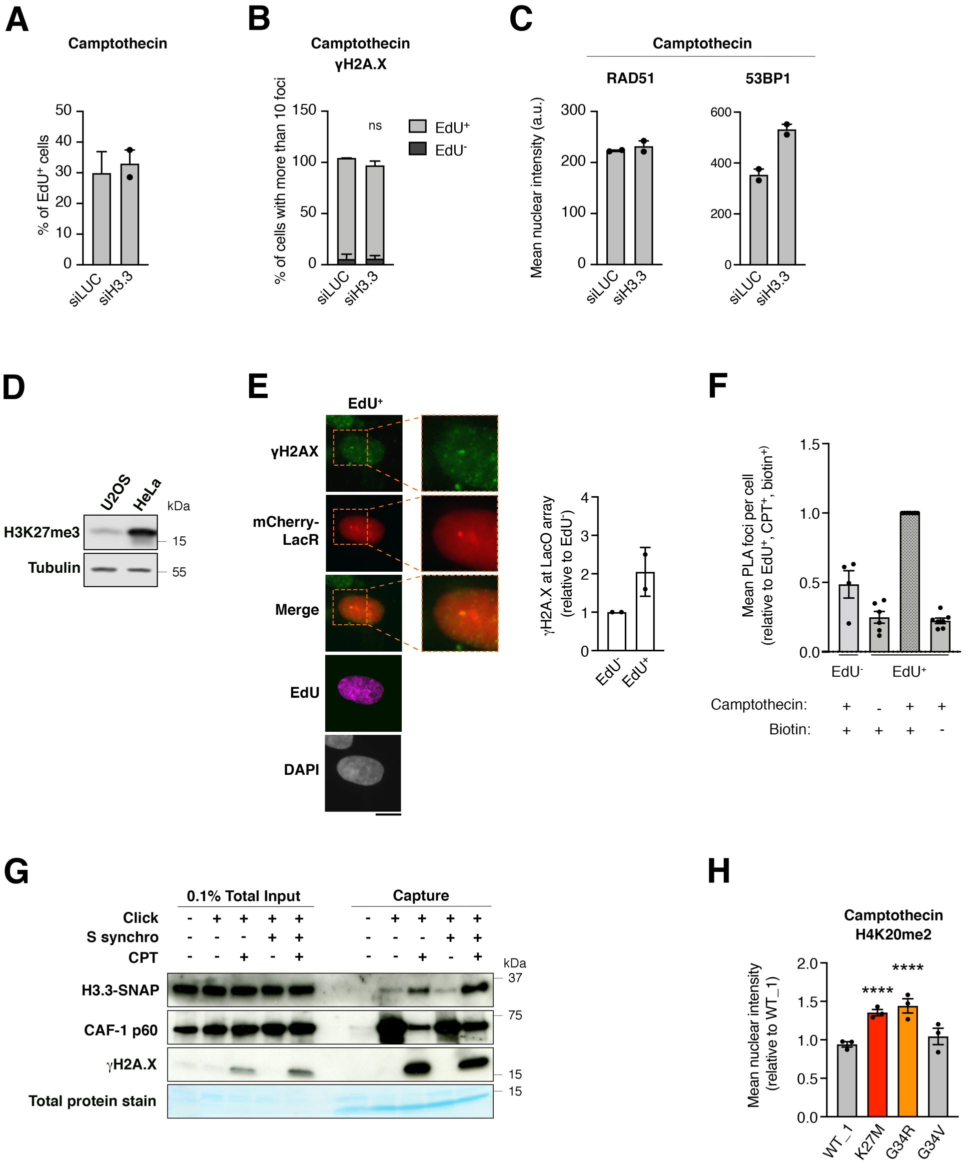


Supplemental Figure S3

**Figure S3. Functional consequences of H3.3 depletion, H3K27 and H4K20 methylation levels and H3.3 deposition at damaged replication forks in the studied cell lines. Related to Figures 2 and 3.** (A-B) Analysis of EdU incorporation (A) or γH2A.X foci formation in $EdU^+$ and $EdU^-$ populations (B) in U2OS cells transfected with siRNAs against Luciferase (siLUC, control) or H3.3 (siH3.3) and treated with camptothecin (3 h, 0.1 μM). Bar graphs depict the percentage of S phase cells ($EdU^+$, A) or γH2A.X foci per cell in $EdU^+$ and $EdU^-$ cell populations (B). (C) Immunofluorescence analysis of nuclear RAD51 or 53BP1 protein levels (a. u., arbitrary units) in U2OS cells transfected with siRNAs against Luciferase (siLUC, control) or H3.3 (siH3.3). (D) Western blot analysis of H3K27me3 levels in U2OS and HeLa cell lines (Tubulin, loading control). (E) Quantification of γH2A.X accumulation at LacR-occupied LacO array analyzed by immunofluorescence in $EdU^+$ and $EdU^-$ U2OS cells stably expressing SNAP-tagged H3.3 wild-type and transfected with mCherry-Lac repressor (LacR). Scale bar, 10 μm. Mean ± SEM from two independent experiments, n>20 per sample for each experiment. (F) Quantification of SNAP-PLA colocalization foci between new H3.3 and γH2A.X in $EdU^+$ and $EdU^-$ U2OS cells stably expressing wild-type H3.3, untreated or treated with camptothecin (3 h, 0.1 μM). A biotin-free sample is included as negative control. Mean ± SEM from up to seven independent experiments, with n>130 per sample for each experiment. (G) Western blot analysis of input and capture samples from iPOND experiments performed in U2OS cells stably expressing wild-type H3.3-SNAP, asynchronous or synchronized in S phase, untreated or treated with camptothecin (3 h, 1 μM). Click -, negative control (no biotin). Detection of the replication-coupled chromatin assembly factor CAF-1 is used as a positive control for undamaged RFs and the DNA damage marker γH2A.X is used as a positive control of CPT damage. Total protein stain shows the position of the streptavidin monomer, detectable at similar levels in all capture samples. (H) Immunofluorescence analysis of nuclear H4K20me2 protein levels in U2OS cells stably expressing wild-type (WT_1) or mutant H3.3 and treated with camptothecin (3 h, 0.1 μM). Intensity is expressed as relative to WT_1. Mean ± SEM from three or two (C) independent experiments, with n>119 per sample for each experiment. Statistical significance is calculated by two-way ANOVA (B) or one-way ANOVA (H). *: $p< 0.05$; **: $p< 0.01$; ***: $p< 0.001$; ns: $p> 0.05$.

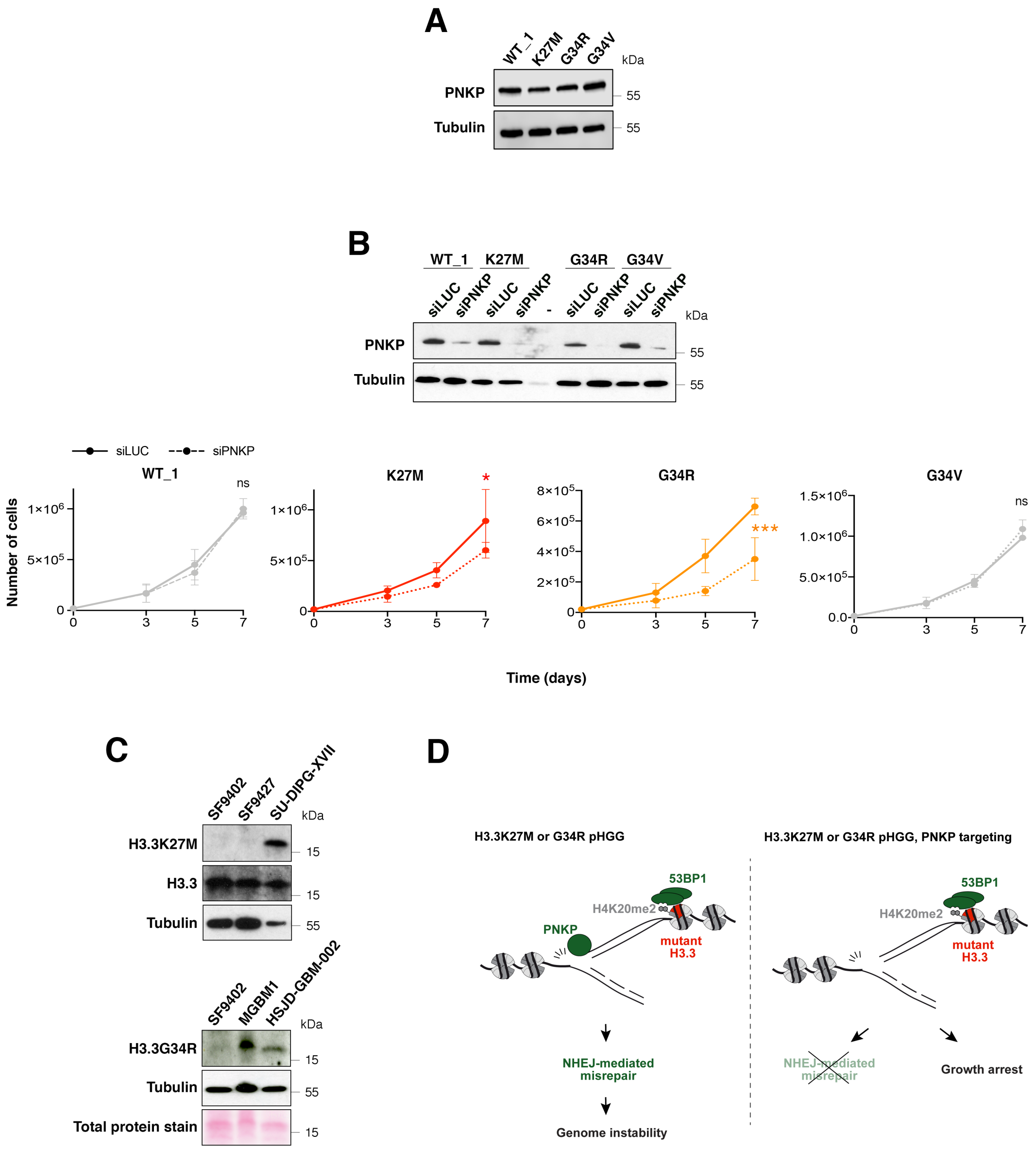


Supplemental Figure S4

**Figure S4. Targeting PNKP in H3.3K27M and G34R mutant cells. Related to Figure 4.** (A) Western blot analysis of whole cell extracts showing PNKP protein levels in U2OS stable cell lines expressing wild-type H3.3 (WT_1) or the indicated mutants. (B) Western blot analysis and proliferation assays in U2OS cells stably expressing wildtype H3.3 (WT_1) or the indicated mutants and transfected with siRNAs against Luciferase (siLUC, control) or PNKP (siPNKP). The western blot shows siRNA efficiencies. (C) Western blot analysis depicting endogenous H3.3 mutations (H3.3K27M and G34R) in a panel of patient-derived pHGG cell lines (Tubulin and total protein stain, loading controls). (D) Current model depicting PNKP-mediated misrepair of S phase DNA damage in K27MH3.3 and G34R (left) and the consequences of PNKP targeting in these cells (right). Statistical significance is calculated by non-linear regression analysis with a polynomial quadratic model (B). *: $p < 0.05$; **: $p < 0.01$; ***: $p < 0.001$; ns: $p > 0.05$.

**Table S1. Proximity interactome of wild-type H3.3 vs. H3.3 K27M or H3.3 G34R mutants. Related to Figure 4.** Proteins associated with wild-type (WT) and mutant H3.3 (K27M, G34R) identified by proximity-dependent biotinylation (BioID) in HEK293 T-REx cells expressing BirA*-tagged H3.3 proteins. Significant hits are highlighted in colors (blue, depleted in the mutant compared to wild-type; red, enriched). Common interactors between H3.3K27M and G34R are highlighted in dark colors (common hits tab).
